# Elevator-type Mechanism of the Cyanobacterial Bicarbonate Transporter

**DOI:** 10.1101/2022.06.08.495363

**Authors:** Matthew C. Chan, Yazeed Alfawaz, Arnav Paul, Diwakar Shukla

## Abstract

Cyanobacteria are responsible for up to 80% of aquatic carbon dioxide fixation and have evolved specialized carbon concentrating mechanism to increase photosynthetic yield. As such, cyanobacteria are attractive targets for synthetic biology and engineering approaches to address the demands of global energy security, food production, and climate change for an increasing world’s population. The bicarbonate transporter BicA is a sodium-dependent, low-affinity, high-flux bicarbonate symporter expressed in the plasma membrane of cyanobacteria. Despite extensive biochemical characterization of BicA, including the resolution of the BicA crystal structure, the dynamic understanding of the bicarbonate transport mechanism remains elusive. To this end, we have collected over 1 ms of all-atom molecular dynamics simulation data of the BicA dimer to elucidate the structural rearrangements involved in the substrate transport process. We further characterized the energetics of the cooperativity between BicA protomers and investigated potential mutations that are shown to decrease the free energy barrier of conformational transitions. In all, our study illuminates a detailed mechanistic understanding of the conformational dynamics of bicarbonate transporters and provide atomistic insights to engineering these transporters for enhanced photosynthetic production.

## Introduction

Marine cyanobacteria, also known as green-blue algae, are estimated to contribute at least 30-80% of the Earth’s total primary production.^1,2^ In aqueous solutions, carbon dioxide (CO_2_) readily interconverts between carbonic acid (H_2_CO_3_) and bicarbonate ions (HCO_3_ ^−^). Unlike CO_2_, HCO_3_ ^−^ cannot freely diffuse through the plasma membrane and thus requires specialized integral membrane transporters to accumulate inorganic carbon for photosynthesis and carbohydrate production. Three bicarbonate transporters have been identified to be ubiquitously expressed in the cyanobacteria plasma membrane: BicA, a sodium-dependent, high-flux, low-affinity bicarbonate symporter; SbtA, a sodium-dependent, high-affinity symporter, and BCT1, a four-subunit bicarbonate transporter belonging to the ATP-binding cassette family^3,4^ (Figure 1A). To date, the solved structure of BicA^5^ (Figure 1B) and most recently of SbtA^6^ have illuminated the molecular architecture of the overall topology and substrate binding site among these critical transporters.

**Figure 1:**
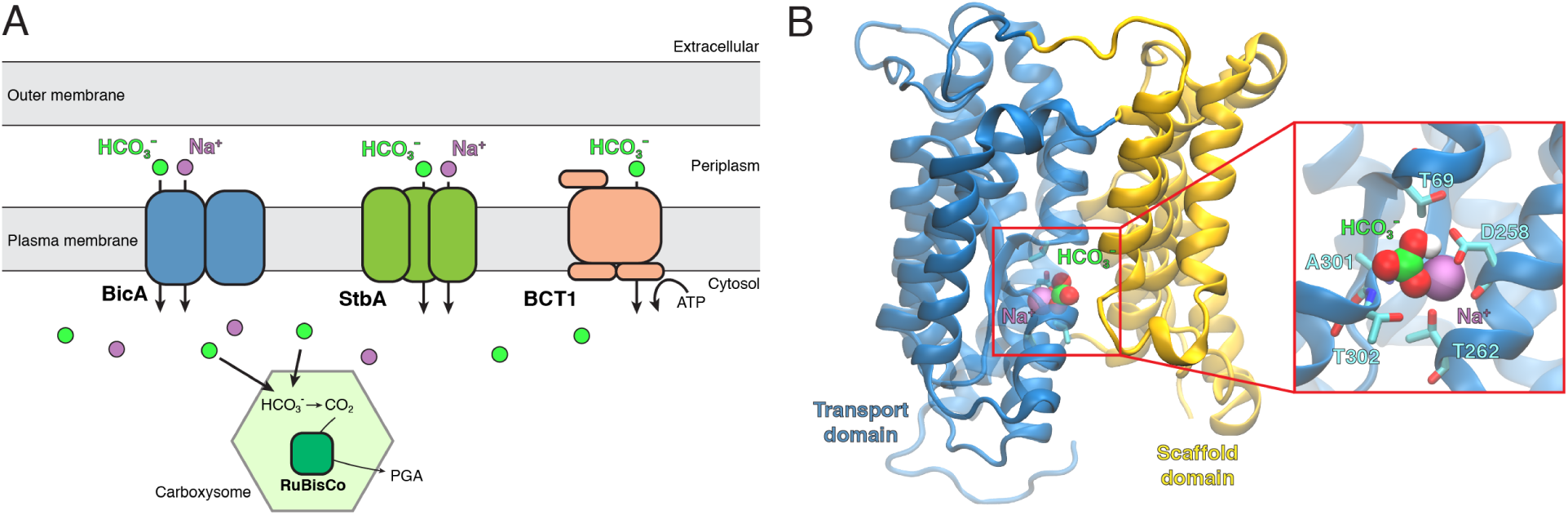
Cyanobacteria bicarbonate uptake transporters. (A) Schematic of select bicarbonate transporters expressed in the cyanobacteria plasma membrane. Transporters are depicted as follows, BicA, a sodium-dependent dimer, blue; StbA, a sodium-dependent trimer, green; BCT1, a four-subunit ATP-binding cassette transporter. Bicarbonate anions that are transported into the cytosol are then concentrated in the carboxysome, converted to carbon dioxide via carbonic anhydrase, and finally undergo photorespiration to form phosphoglyceric acid (PGA) via RuBisCo. (B) MD equilibrated structure of the BicA based on the crystal structure PDB: 6KI1. The cytoplasmic STAS domain is not shown for clarity. The transport domain and scaffold domain are colored as blue and yellow, respectively. The bound substrates are represented as spheres. Residues that coordinate the binding of the substrates are shown as sticks.

In C3 crops, which include rice, barley, and wheat, carbon fixation via RuBisCo (ribulose-1,5-bisphosphate carboxylase/oxygenase) is notoriously known to be inefficient.^7^ As such, a possible approach to increases crop yield is to incorporate the efficent CO_2_-capturing mechanisms utilized by cyanobacteria^8^ into crops. Inorganic carbon transporters are one of two components that make up an effective CO_2_-capturing mechanism, the other being the carboxysomes, which are specialized protein micro-compartments that house RuBisCo and carbonic anhydrase to concentrate CO_2_ for efficient carbon fixation.^9^ Kinetic modeling has proposed that introducing cyanobacteria bicarbonate transporters to the chloroplast of C3 crops may enhance photosynthetic yield by *∼*10%, while adding the carboxysome system may further increase yield as much as *∼*60%.^10^ Incorporating either bicarbonate transporters and the carboxysome involved the synthetic addition of foreign genes to the chloroplast of plastid genome. However, whereas bicarbonate transporters are simply encoded as single genes, the *in vivo* assembly of the carboxysome requires multiple proteins and presents inherent difficulties to simultaneously introduce all the required genes. As such, bicarbonate transporters are attractive candidates for engineering terrestrial crops to enhance inorganic carbon accumulation.^11,12^ Additionally, increased carbon availability promotes cyanobacteria growth which may be used for the production of biofuels and other bioproducts.^13^

The cyanobacteria bicarbonate transporter BicA is a member of the solute carrier 26 (SLC26/SulP) family of anion transporters. Members of this family contain an N-terminal transmembrane (TM) domain comprised of 14 helices arranged in a 7+7 inverted repeat topology and a cytoplasmic C-terminal domain known as the sulfate transporter and anti-sigma factor antagonist, or STAS, domain.^5^ Despite low sequence conservation, transporters in the SLC4 and SLC23 families share the similar 7+7 transmembrane architecture, but most notably lack the STAS domain.^14^ Furthermore, SLC26 transporters have been shown to adopt a unique dimer interface that involves TM helices 13 and 14, whereas TM helix 6 forms the dimer interface for SLC4 transporters, and TM helices 5 and 12 for SLC23.^15–17^ Biophysical, structural, and computational studies^18–20^ have illuminated the SLC26 family and similar related families to adopt a canonical alternating-access model in which the transporter undergoes a series of structural rearrangements to enable access of an orthosteric substrate binding site from either the extracellular or intracellular side. ^21^ More specifically, the mode of transport of SLC26 transporters has been proposed to be an elevator-like mechanism, in which helices 1-4 and 8-11 form a mobile transport domain that translates across the membrane, thereby transporting substrates in and out of the cell (Figure 1B). TM helices 5-7 and 12-14 form the scaffold domain that remains rigid and is primarily involved in oligomeric assembly. Analogous SLC26 transporters in humans are involved in the exchange of anions throughout the body and mutations in these transporters are associated with various disorders such as cystic fibrous, chloride diarrhea, and chondrodysplasia. ^22^

It is estimated that by 2050, the global food production must be doubled in order to sustain a growing population.^23,24^ In order to address the concern of global food security and sustainable energy, understanding the molecular mechanism of bicarbonate transport in cyanobacteria may serve as the basis for enhancing the efficiency of crop yield and biofuel production. While the resolved structure of BicA provides invaluable structural information, the conformational dynamics and energetics involved in the substrate translocation process may not be elucidated from a single structure and therefore remain elusive. With the recent surge in the computational efficiency of graphical processing units and numerical algorithms, molecular dynamics (MD) simulations combined with Markov state modeling present a robust approach to characterize complex protein dynamics at atomistic resolution.^25,26^ Recent efforts in Markov state modeling have characterized the conformational heterogeneity of proteins of key interest to the plant biology community including phytohormone receptors,^27–29^ and circadian clock photoreceptors.^30–32^ Several membrane transporters have also been investigated using these methodologies including sugar transporters (SWEETs and SemiSWEETs),^33–35^ bacterial nitrate transporters,^36^ human neurotransmitter^26,37^ and peptide transporters.^38^ However, these transporters follow either the rocker-switch or rocking bundle mechanisms of alternate-access to facilitate the substrate transport. ^39,40^ BicA is distinct from these transporters because it follows the elevator-type mechanism,^41–46^ where the transport domain undergoes a translation relative to the scaffold domain to achieve alternate-access required for substrate transport. ^5^

In this current study, we employed long-timescale all-atom MD simulations to provide a fully atomistic and dynamic perspective into the bicarbonate transport mechanism of BicA. We further analyzed the simulation dataset using Markov state modeling^26^ to quantify the thermodynamics of the elevator-like mechanism and its associated structural rearrangements. Finally, we investigated the effects of BicA mutations on the transporter structure and dynamics and present a mechanistic basis for mutations that may be introduced to BicA to enhance bicarbonate transport activity. Overall, our computational study provides an atomistic level perspective into the molecular mechanisms of BicA bicarbonate transport which may be used for further engineering of cyanobacteria and plants.

## Methods

### MD simulations of pure cyanobacteria plasma membrane

To characterize the structural dynamics of BicA, we first sought to model a physiological membrane environment. Based on previous experimental characterization of cyanobacteria membranes, we constructed a symmetric lipid membrane containing the monogalactosyl diacylglycerol (MGDG), digalactosyl diacylglycerol (DGDG), sulfoquinovosyl diacylglycerol (SQDG), and phosphatidylglycerol (PG). The total number of each lipid species and lipid tail saturation are detailed in Table S1. A total of 130 lipid molecules per leaflet were assembled using PACKMOL.^47^ Water molecules and sodium ions to neutrailze the system were further added. In all, the final MD membrane system contained 152 MGDG molecules, 42 DGDG molecules, 42 SQDG molecules, 24 PG molecules, 152 sodium ions, and 19,998 water molecules totaling 140,834 atoms in a rectilinear box of 140 x 101 x 105 Å^3^. A total of three membrane systems, randomizing the initial lipid placement, were constructed.

The MD systems were parameterized using the CHARMM36m force field. The parameters for the saturated lipids tails (18:3*γ*/16:0, 18:2/16:0, 18:3*α*/16:0), which are not originally parameterized in CHARMM36, were derived analogous from parameters of related lipid molecules in the CHARMM36 molecule set. The *psf* topology and coordinating file were created using the VMD psfgen plugin and converted to AMBER *prmtop* topology and *rst7* coordinate files using the *chamber* module of the ParmEd package.

Simulations were performed on the AMBER18 package using the *pmemd* GPU acceleated module. The MD system was first minimized 7,000 steps using the steepest descent method followed by 93,000 steps using the conjugate gradient method. Prior to production simulations, the system was heated to 300K in 100K increments for 1 ns each while restraining the lipid head group atoms with a force constant of 1 kcal/mol-Å^2^. Production simulations were performed in an NPT ensemble using Langevin dynamics with a damping coefficient of 1 ps^-1^ at 300K, 1 bar, and positional restraints removed. A Monte Carlo barostat with an update interval of 100 steps was used to maintain pressure. A 12 Å distance cutoff was applied to calculate nonbonded interactions. Long-range electrostatic interactions were treated with the Particle mesh Ewald method. Hydrogen bonds were constrained using the SHAKE algorithm. An integration timestep of 2 fs was used for membrane simulations. Each MD replicate was simulated for 250 ns with a trajectory frame saving rate of 100 ps.

We noted that a previous computational study identified errors in using a similar parameter combination of the Monte Carlo barostat and 12Å hard cutoff.^48^ We have compared the dynamics of our lipid membrane with a 12Å cutoff/10Å switch versus just a 12Å cutoff/no switch function. In both simulated setups, the membrane approaches equilibrium after *∼*50 ns (Figure S1). For the 12Å cutoff, we calculated the average membrane thickness to be 37.2 ± 0.3 Å and area-per-lipid (APL) of 58.8 ± 0.5 Å^2^ (average *±* standard deviation, over last 50 ns). In comparison, the 12Å cutoff/10Å switch resulted in an average membrane thickness of 36.5 ± 0.3 Å an APL of 61.4 ± 0.5 Å^2^. Overall, we find that systems remain stable throughout the simulations, with small differences in the APL and membrane thickness between the two simulation parameters.

### Modeling of the full-length BicA dimer system

The three-dimensional coordinates of the resolved BicA crystal structure (PDB:6KI1, 6KI2)^5^ were used as the starting structure for simulations. First, transmembrane helix 14 was modelled based on the SLC26Dg structure (PDB: 5DA0) To model the full-length BicA dimer, two transmembrane domains were aligned to the SLC26a9 murine transporter, ^16^ in which transmembrane helices 13 and 14 formed the dimeric interface.^17^ The STAS domain was placed under the transmembrane domain and residues that linked the two domains were modelled with MODELLER.^49^ We found that the resulting full-length BicA dimer to modestly fit in the cryo-EM map,^5^ which may be attributed to its low-resolution or reconstruction in detergent which may impact the packing of membrane proteins. The modeled dimer, containing residues 2-547, was embedded in the cyanobacteria plasma membrane and solvated with TIP3P water molecules. 10 bicarbonate anions were randomly placed in the solvent and sodium ions were added to neutralize the system. Protonation states were assigned based on pKa calculations from PROPKA3.4**^?^** and a system pH of 8.^5^ The final BicA dimer system consisted of 2 BicA protomers, 152 MGDG molecules, 42 DGDG molecules, 42 SQDG molecules, 24 PG molecules, 10 bicarbonate anions, 80 sodium ions, and 30,200 water molecules totaling in 141,886 atoms in a rectilinear box of 140.0 x 101.0 x 126.0 Å^3^.

The alternative structure prediction of the BicA dimer was generated using AlphaFold v2.2.0 in tandem with the multimer mode.^50,51^ The AlphaFold prediction was performed using the following parameters: `--max template date=2022-05-01, model preset=multimer, --norun relax, --db preset=reduced dbs`. The output structure from AlphaFold was not used for simulations in this study.

### MD simulations of full-length BicA dimer

The BicA dimer system was parameterized using the CHARMM36m force field and conducted on the AMBER18 package. Prior to production simulation, the system was minimized for 5,000 steps using the steepest descent method, followed by 45,000 steps using the conjugate gradient method. The BicA dimer was then subjected to 10 pre-production simulations with varying atoms restrained, totaling in 150 ns. A detailed list of temperatures, restrained atoms, and simulation length for each pre-production step is presented in Table S1.

Production simulations were performed under the same conditions and parameters as the previous described pure membrane system, with the exception of the use of a 4 fs integration timestep and hydrogen mass repartition.^52^ To determine the stability of the full-length BicA dimer, a total of 7 simulations were initiated from the last pre-production step and simulated for 700 ns. The resulting trajectories from these simulations yielded BicA to remain in the inward-facing state. As such, to explore the conformational space of BicA, we adaptively sampled the conformational landscape based on the distances of the substrate to the binding sites and the displacement of the transport domain.^26^ The adaptive sampling scheme involves iteratively performing many independent simulations in parallel, followed by clustering the simulation data based on a geometric metric. The subsequent round of adaptive sampling simulations is seeded from the least populated clusters to maximize the probability of observing new conformations. However, after 14 rounds of adaptive sampling, we did not observe either BicA protomer to transition from the inward-facing state (Figure S2). As such, we seeded subsequent rounds from a targeted MD trajectory in which simulated the transition from inward-facing to outward-facing (Figure S6). The targeted MD simulation was performed using NAMD2.14^53^ using a 200 kcal/mol-Å^2^ force constant and an outward-facing/outward-facing BicA homology model based on the NBCe1 cryo-EM structure (PDB: 6CAA)^54^ as the target structure. A total of 29 adaptive rounds, in which individual trajectories were 60 ns long, were conducted and totaled in 1.003 ms of aggregate simulation data (Table S2). Production simulations were performed on the Blue Waters supercomputer on Tesla K20 GPUs.

### Trajectory analysis

Trajectories were processed with in-house scripts utilizing the CPPTRAJ, pytraj,^55^ and MDTraj^56^ packages. Simulation trajectories were visualized using Visual Molecular Dynamics (VMD) 1.9.3.^57^ Residue contact probabilities were calculated using the GetContacts (https://getcontacts.github.io/) python package. Cross-correlation analysis of BicA was performed using the Bio3D R package.^58^

### Markov state modeling

All 1.003 ms of aggregate simulation data from adaptive sampling simulations were used to construct a reversible Markov state model (MSM) using the pyEMMA python package.^59^ The simulation data were featurized based on the distances of residue pairs that were identified to uniquely form in the inward- and outward-facing states and determined using a residue-residue contact score (RRCS).^60^ In all, 192 distances were identified between the two BicA protomers (Table S3). Additionally, the *z*-components of Asp258 and the transported bicarbonate anion were included as features for the MSM, totaling in 196 cartesian features. The number of time-independent components (tICs) and number of clusters was optimized using a grid search to maximize the VAMP1 score (Figure S3A). The best-scoring model was achieved with 10 tICs and 400 clusters. A lagtime of 10 ns was determined based on convergence of the implied timescales (Figure S3B). The standard error of the free energy landscape was calculated using the bootstrap method by randomly selecting 80% of the trajectories for 500 independent sample (Figure S4). The reversibility of the MSM was assessed by comparing *π_i_P_ij_*= *π_j_P_ji_*and the differences between *π_i_P_ij_*and *π_j_P_ji_*were within 10^-15^ (Figure S5).

### Umbrella sampling simulations

To investigate the energetics of conformational transitions of BicA mutants, we employed umbrella sampling simulations. The displacement between the *z*-component of the transport domain center of mass and the *z*-component of the scaffold domain center of mass was used as the reaction coordinate to describe the conformational transitions from inward-facing to outward-facing. Based on the conformational free energy landscape, one protomer of BicA is predicted to undergo transitions at once, and as such, we employed the umbrella sampling protocol on only BicA monomer A. Structures were drawn from the conformational landscape. A total of 26 windows from *z*-coordinates −8.0 to +4.5, in 0.5Å intervals were used to seed umbrella sampling simulations. Each window was simulated for 9 ns. Umbrella sampling simulations were performed using NAMD2.14^53^ using a 2 fs integration timestep, 12Å cutoff with a 10Å switching distance, and a harmonic force constant of 15 kcal/mol-Å^2^. Simulation frames were saved every 10 ps. Potentials of mean force (PMFs) were calculated using the multistate Bennett acceptance ratio as implimented by the pyMBAR python package.^61^ Secondary structure, computed by DSSP, remained intact throughout all umbrella windows (Figure S7). Convergence and overlap matrices of umbrella sampling simulations are shown in (Figure S8-S13).

### Generation of multiple sequence alignments

Multiple sequence alignments were generated using the ConSurf web server. ^62^ The sequences of BicA, UraA (PDB:5XLS),^18^ and NCBe1 (PBD:6CAA)^54^ were used as input to represent SLC26, SLC23, and SLC4 families. A 95% maximal identity between sequences and a 35% minimal identity between homologs was used to created the alignment. 300 represented sequences were sampled from the list of accepted homologs. Sequence logo figures were generated using the WebLogo server.^63^

## Results and Discussion

### Structural characterization of a full-length BicA dimer in a cyanobacteria plasma membrane

We sought to simulate a full-length model of BicA in a lipid membrane that best resembles the physiological plasma membrane of cyanobacteria. In addition to portraying a realistic molecular environment, the increased membrane complexity may also affect thermodynamics barriers across the conformational landscape.^35^ To this end, we constructed a lipid membrane based on previously characterized compositions determined for cyanobacteria (Figure 2A).^64,65^ Notably, the cyanobacteria plasma membrane is comprised of mainly glycolipids. The constructed membrane consisted of the four unique and most abundant identified lipids: monogalactosyl diacylglycerol (MGDG), digalactosyl diacylglycerol (DGDG), sulfoquinovosyl diacylglycerol (SQDG), and phosphatidylglycerol (PG). The saturation of fatty acid tails were also modeled in accordance to Murata *et al*. ^64^ Further details of the composition of the simulated membrane are listed in Table S1.

**Figure 2:**
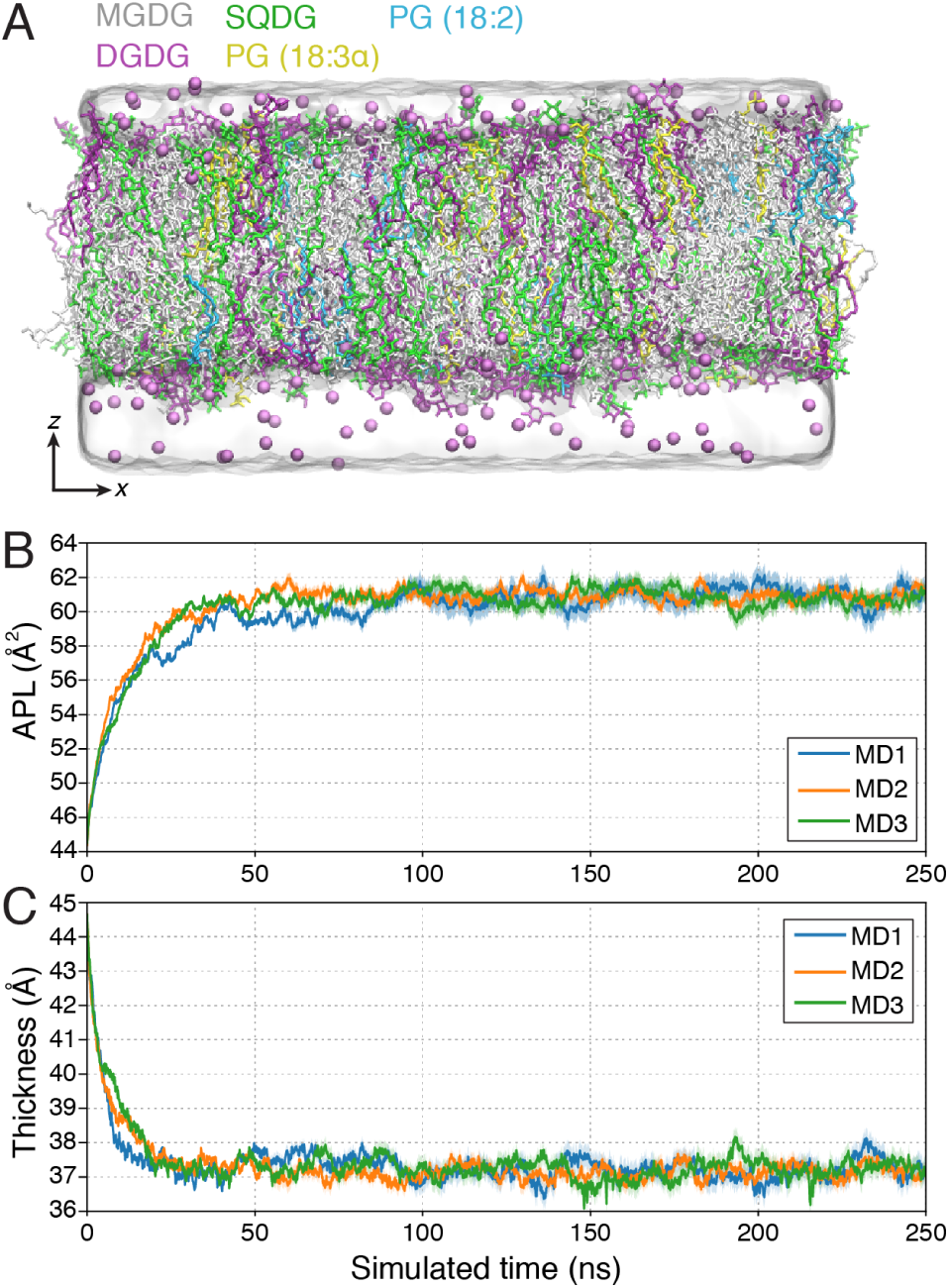
Molecular dynamics simulations of the cyanobacteria plasma membrane. (A) MD snapshot of the simulated cyanobacteria plasma membrane. Lipid molecules are shown as sticks and colored by individual lipid species. Sodium ions are represented as purple spheres. (B, C) Time-resolved measurements (B, area-per-lipid (APL) and C, membrane thickness) of the cyanobacteria lipid membrane. The three MD replicates are colored accordingly. Error bars represented accumulated standard deviation after the initial 50 ns.

To assess the dynamics of the cyanobacterial plasma membrane, all-atom MD simulations were performed using the AMBER18 engine^66^ employing CHARMM36 force fields.^67^ Three independent replicates with unique initial lipid placement were constructed and simulated for 250 ns each. The simulated membrane readily approaches equilibrium after *∼*50 ns, with an average membrane thickness of 37.2 *±* 0.3 Å and area-per-lipid of 58.8 *±* 0.5 Å^2^ (average *±* standard deviation, over last 50 ns) (Figure 2B, C). We find that the physical properties of our simulated membrane is in agreement with the compositionally-similar cyanobacteria thylakoid membrane, previous characterized by simulation.^68^

**Table S1:** Composition for pure cyanobacteria plasma membrane simulations.

| Lipid | Saturation (sn1/sn2) | Number of lipids per leaflet | Percentage |
| --- | --- | --- | --- |
| MGDG | 18:3γ/16:0 | 76 | 58% |
| DGDG | 18:3γ/16:0 | 21 | 16% |
| SQDG | 18:2/16:0 | 21 | 16% |
| PG | 18:2/16:0 | 6 | 5% |
| PG | 18:3α/16:0 | 6 | 5% |

With the cyanobacteria plasma membrane established, we set to construct a full-length dimeric BicA system, using the resolved crystal structures of the transmembrane domain (PDB: 6KI1) and the STAS domain (PDB: 6KI2).^5^ Based on pulsed electron-electron double resonance spectroscopy^17^ and the cryo-EM structure of the dimeric SLC26a9 murine transporter,^16^ the initial orientation of the two BicA protomers was placed where helices 13 and 14 formed the interface. We alternatively modeled a full-length BicA dimer structure using AlphaFold^50^ and observed a similar orientation of the transmembrane domains and its interface (Figure S14). However, the structure predicted by AlphaFold did not model the STAS domain in accordance to the crystal structure or cryo-EM density (Figure S14). As such, we proceeded with the BicA dimer structure based on the two available crystal structures and superposition of other SLC26 transporters. The full-length BicA dimer was embedded in the cyanobacteria plasma membrane and a total of seven MD replicates of 700 ns were performed (Figure 3A). During the pre-production stages of the simulations, we observed the one sodium ion to bind to one BicA monomer and remained bound throughout the 700 ns simulation across the seven replicates. The binding of a sodium ion to the remaining BicA monomer was observed within 200-500 ns of simulation (Figure S15). In both cases, the sodium ion is coordinated by the side chains of Asp258, Thr262, and Thr302, consistent with the resolved crystal structure and previous mutagenesis characterization. ^5^ The bicarbonate anion was not observed to bind in the substrate cavity within the equilibration timescales.

**Figure 3:**
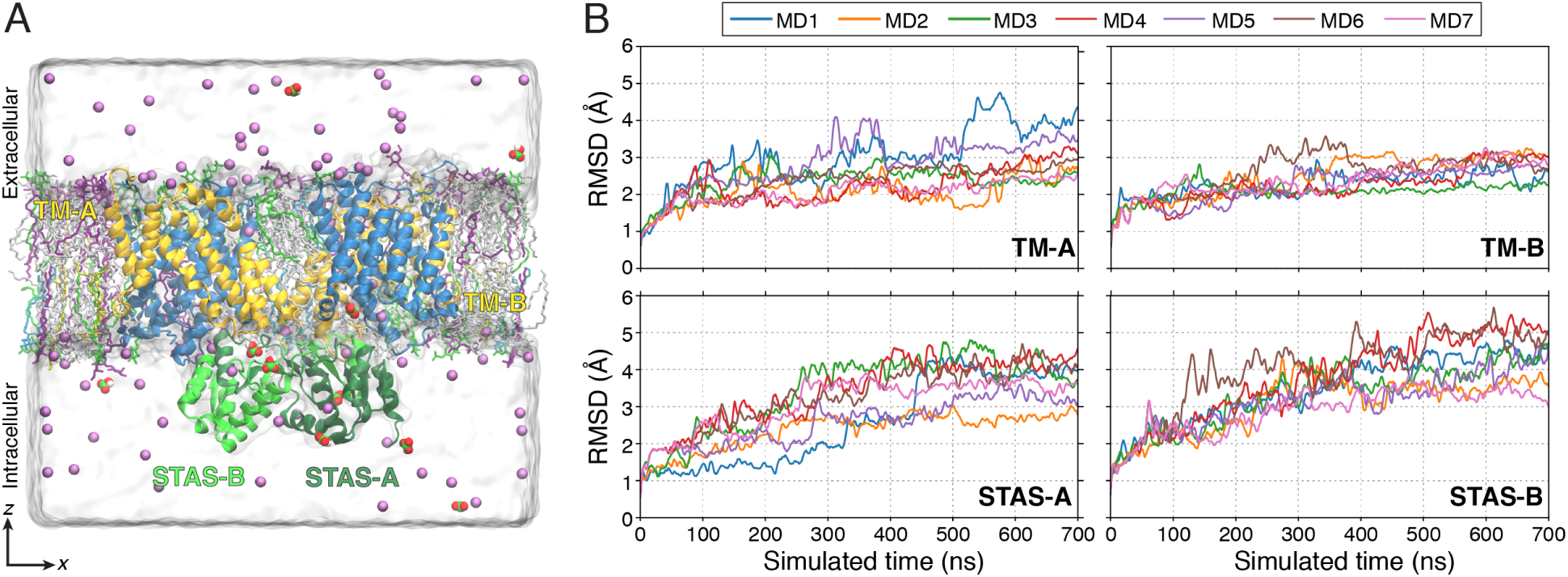
Stability of the full-length BicA dimer model. (A) MD snapshot of the full-length BicA dimer embedded in the cyanobacteria plasma membrane. Lipid molecules are shown as sticks and colored by lipid species as shown in Figure 2A. The BicA dimer is shown in cartoon representation and colored as follows: yellow: blue domain; blue: transport domain; green: STAS domain. Individual BicA protomers are labeled as A and B. Sodium ions are shown as purple spheres. Bicarbonate ions are shown as red and green spheres. (B) Time-resolved root mean squared deviations (RMSD) of individual BicA transmembrane (TM) and STAS domains across the 7 MD simulations. RMSD was calculated based on the starting structure of the 700 ns simulation. Individual MD replicates are colored accordingly.

The simulations reveal that the transmembrane domains of BicA remain relatively stable (C*α* RMSD *<* 3.5 Å), whereas the cytoplasmic STAS domain deviated from its initial structure (C*α* RMSD *>* 3.5 Å) (Figure 3B). Moreover, upon equilibrating the full-length BicA system, we observed particularly the *α*2 helices to unwind and propagate the collapse of the domain. Indeed, the structural elements that form the STAS domain architecture are not maintained throughout the 700 ns long simulations (Figure S16). The inherent differences between the simulated and experimentally determined structure may be attributed to artificial crystal contacts formed during crystallography or the lack of the transmembrane domains being coexpressed to mediate the folding of the STAS domain. Functionally, the STAS domain of the sulfate transporter Sultr1;2 in *Arabidopsis thaliana* has been characterized to interact with cysteine synthase (O-acetylserine (thiol)lyase) to regulate the transporter function and mediate the cellular sulfur concentration.^69^ The association of the STAS domain with other regulatory proteins is further exhibited in SLC26A3 transporter in humans with implications to cystic fibrosis. ^70^ As such, it is likely the STAS domain may adopt various conformations in solution and be stabilized upon association. We further simulated a BicA dimer with the STAS domains removed and did not observe difference in dimer stability of the transmembrane domain (Figure S17).

### Structural requirements and energetics of BicA conformational transitions

As the timescales of large structural rearrangements and substrate transport may occur on the orders of microseconds or greater, ^71,72^ observing these long timescale processes through conventional MD approaches may present inherent challenges in achieving adequate conformational sampling. As such, to simulate the bicarbonate transport process of BicA, we implemented a Markov state model (MSM) based adaptive sampling scheme to maximize the exploration of the conformational landscape.^26^ In brief, the adaptive sampling protocol is an iterative approach in which multiple simulations are conducted in parallel and then clustered using a K-means algorithm based on geometric criteria. To sample the BicA substrate translocation process, the distances between substrates and binding site and the *z*-component of the transport domain were chosen as the adaptive sampling metrics. To maximize the likelihood of exploring new conformations, structures from the least populated states are seeded for the subsequent round of simulation. Furthermore, to expedite the sampling, we seeded simulations from a targeted MD trajectory that captured the transition from inward-facing to outward-facing (Figure S6). A total of 1.003 millisecond of aggregate simulation data were collected and used to construct a MSM.^73^ We note that the conformational sampling performed for this study simulates the export of bicarbonate to the extracellular side. Though BicA is responsible for concentrating inorganic carbon in the cell, the benefit of the Markov state modeling is representing the transport process as a reversible process and calculating the reversible transition probabilities between states, thereby capturing the bicarbonate import process.

By projecting the MSM-weighted simulation data on the reaction coordinates defined by the *z*-component of the Asp258 C*α* atom (the residue that coordinates binding of the bicarbonate anion, Figure 1B), the conformational free energy landscape illustrates the cooperativity of the two BicA protomers (Figure 4A). Inward-facing conformations, in which the substrate binding site is accessible from the intracellular solvent (Figure 4B), are energetically stable with a relative free energy of 0-1 kcal/mol. Furthermore, the simulations reveal that the BicA protomer may independently undergo structural rearrangements to form outwardfacing states in which the transport domain has shifted *∼*6Å and the substrate binding site is now accessible to the extracellular space (Figure 4B). Similar to other transporters that adopt the elevator mechanism,^74,75^ the BicA transport domain simultaneously undergoes rotational and translational motion to fulfill alternate access (Figure S20).

**Figure 4:**
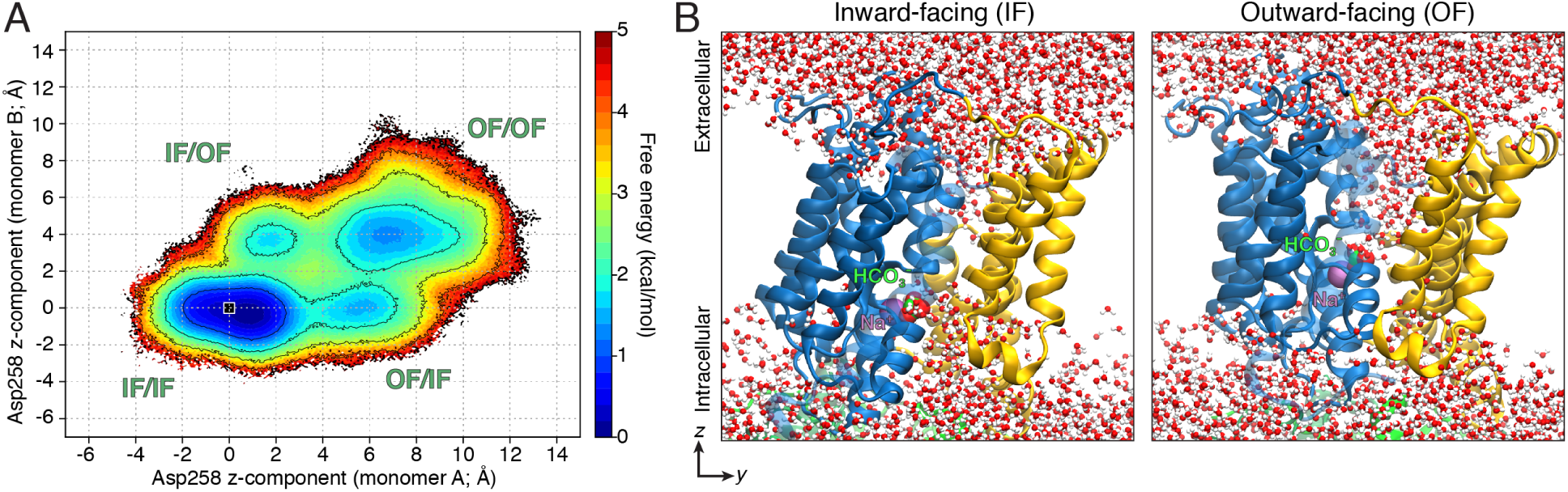
Energetics of BicA conformational transitions. (A) MSM-weighted conformational free energy landscape of BicA. The simulation data are projected on the axis defined by the *z*-component of the C*α* atom of Asp258 for the respective BicA protomers. The displacement of Asp258 is measured with respected to the initial structure used for adaptive sampling simulations and indicated by the black square. Standard error measurement of the free energy landscape is presented in Figure S4. (B) Representative MD snapshots of BicA showing the solvent accessibility of the substrate binding site in the inward-facing (IF) and outward-facing (OF) conformation. The transport domain is shown as blue cartoon, while the scaffold domain is colored in yellow. Water molecules are shown as red and white spheres. STAS domain not shown for clarity.

The free energy barrier associated with transitions from the inward-facing to outward-facing for a single BicA protomer is estimated to be *∼*2.5-3 kcal/mol (Figure 4A, S18). Based on the sampling seeded from the targeted MD trajectory, structures in which both BicA protomers form the outward-facing conformation are stable with a relative free energy minima of *∼*2 kcal/mol. However, the transition free energy barriers for the remaining BicA protomer to adopt the outward-facing state, given that the other protomer is already outward-facing, is *∼*3-4 kcal/mol. Likewise, for both protomers to simultaneously transition to the outward-facing state is energetically less favored with free energy barriers of *∼*4-5 kcal/mol (Figure S18). We have seeded three simulations of BicA in the OF state and collected 700 ns of free dynamics in a similar manner as the inward-facing simulations. Compared to the IF state simulations, we find similar RMSD deviations from the respective initial state and reveal that the OF conformation remains stable with a RMSD of 4Å when compared to the initial OF state (Figure S19). Our simulations were unable to capture strong coupling effects between the two BicA protomers during IF to OF transitions (Figure S21A), Estimation of the mean first passage time between conformational states identified the timescale for transition on the order of hundreds of microseconds. Specifically, the transition from IF/IF state to OF/OF state directly is the fastest pathway with a MFPT of 217 *µ*s, while transition involving intermediate states (OF/IF or IF/OF) is slower (Figure S21B). Overall, the conformational free energy landscape suggests one protomer of BicA actively undergoes structural transitions in the dimeric state, consistent with previous studies of the SLC26Dg fumarate transporter.^18^

We note the presence of two proline residues, Pro122 and Pro341, that flank the transport domain (Figure 5A). Sequence analysis reveals that in 300 homologs, Pro341 is absolutely conserved whereas Pro122 is substituted for Ser in a few homologs (Figure 5B). As the proline residue adopts a cyclic side chain that uniquely constrains the protein backbone, we hypothesized if the steric effects provide the structural requirements for the conformational transitions of BicA. To investigate the effects of the proline residues on the transport dynamics, we implemented umbrella sampling simulations and calculated the potentials of mean force (PMF) profiles of BicA to transition from inward-facing to outward-facing. The conformational free energy landscape suggest that a single BicA protomer is more favored to transition rather than both simultaneously. As such, umbrella sampling simulations were initiated from MD snapshots of the BicA monomer A obtained from the adaptive sampling simulations. Umbrella sampling simulations were conducted with the NAMD2.14 package.^53^

**Figure 5:**
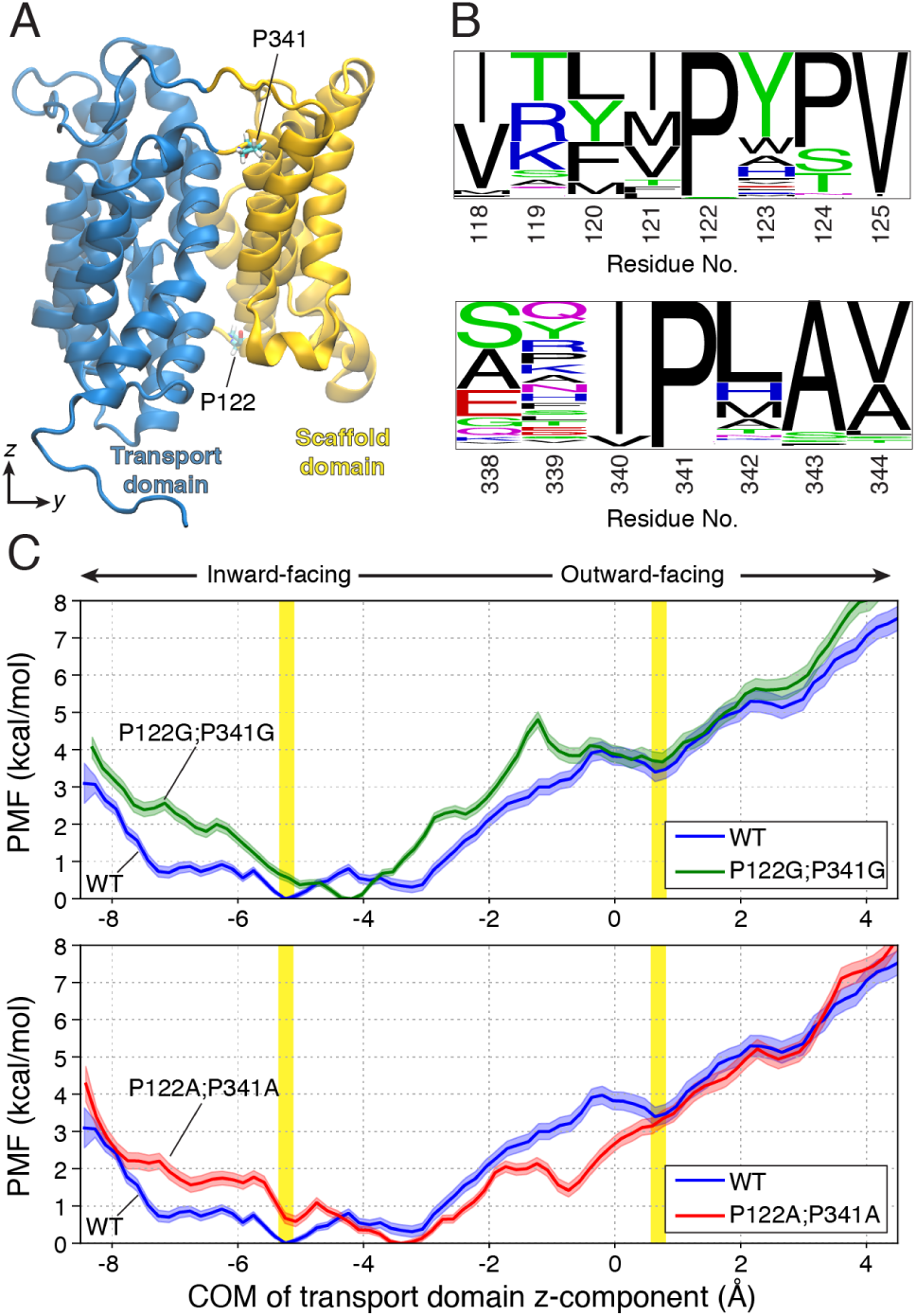
Conserved prolines residues flank the BicA transport domain. (A) Structure of the transmembrane domain of BicA, colored by transport and scaffold domain in blue and yellow respectively. Proline residues, P122 and P341, investigated are indicated and shown as sticks. (B) Sequence logo representation^63^ of the proline and adjacent residues depicting the amino acid frequency of 300 homologs. Size of the amino acid font represents its respective frequency in the multiple sequence alignment. (C) Potentials of mean force (PMF) profiles for inward-facing to outward-facing transitions of wild-type (WT) and mutant BicA. BicA systems are colored as follows, wild-type: blue, P122G;P341G: green, P122A;P341A: red. *x*-axis represents the *z*-coordinate displacement of the center of mass (COM) of the transport domain with respect to the center of mass of the scaffold domain. Inward- and outward-facing conformations are highlighted as vertical yellow bars. Error bars represents standard error.

In the wild-type BicA system, the highest free energy barrier is associated with the transition to the outward-facing state, with a barrier of 3.97 *±* 0.24 kcal/mol. When the Pro122 and Pro341 are mutated to glycine residues (Pro122Gly;Pro341Gly), we observed the stability of the outward-facing state to be similar to that of the wild-type (wild-type: 3.39 *±* 0.25 kcal/mol, Pro122Gly;Pro341Gly: 3.66 *±* 0.24 kcal/mol), but the free energy barrier has now increased to 4.81 *±* 0.21 kcal/mol (Figure 5C). Contrary to prolines, glycine residues provide innate flexibility of the peptide backbone given the lack of a heavy atom side chain. However, such flexibility did not provide necessary structural requirements to promote the formation of the outward-facing state. We further simulated and calculated the PMF profile for the respective alanine mutant (Pro112Ala;Pro341Ala) and observed that the free energy barrier is reduced to 2.14 *±* 0.16 kcal/mol (Figure 5C). Moreover, the Pro112Ala;Pro341Ala mutant stabilizes an intermediate outward-facing state, but the complete outward-facing state remains of similar stability (wild-type: 3.39 *±* 0.25 kcal/mol, Pro122Ala;Pro341Ala: 3.15 *±* 0.21 kcal/mol). As alanine residues enable more structural constraints on the backbone compared to the glycine residues, the PMF profiles suggest that unique dihedral constraints provided by proline residues facilitate the necessary structural rearrangements of the transport domain of BicA, although we cannot comment on how these mutants may affect transporter expression, biogenesis, folding, or stability.

### Hydrophobic interactions mediate closure of the transport domain

Membrane transporters adopt a canonical series of structural rearrangements that facilitate proper substrate transport across the membrane, otherwise known as the alternating access mechanism.^76^ As such, the substrate binding site is accessible to either the intracellular or extracellular space at a given time. Simulations of BicA reveal that the closure of the transporter from either side is facilitated by hydrophobic residues that line the substrate translocation pathway (Figure 6). Specifically, in the inward-facing conformation, closure from the extracellular side is primarily mediated by transmembrane helices 1 and 3 of the transport domain and helices 5, 7, 12, and 14 of the scaffold domain (Figure 6A). Upon substrate transport, as the transport domain shifts across the membrane, intracellular gate is formed by residues on helices 8 and 10 with the scaffold domain (Figure 6B). As per the elevator-like mechanism, the scaffold domain serves as a shared gate between intracellular and extracellular residues. Furthermore, residues that comprise the hydrophobic gate are generally conserved among other transporters that adopt the 7+7 transmembrane helix topology (Figure S22, S23). We expect that the hydrophobic residues in the respective positions of SLC4 and SLC23 transporters to adopt a similar role in regulating the opening and closure of the transporter.

**Figure 6:**
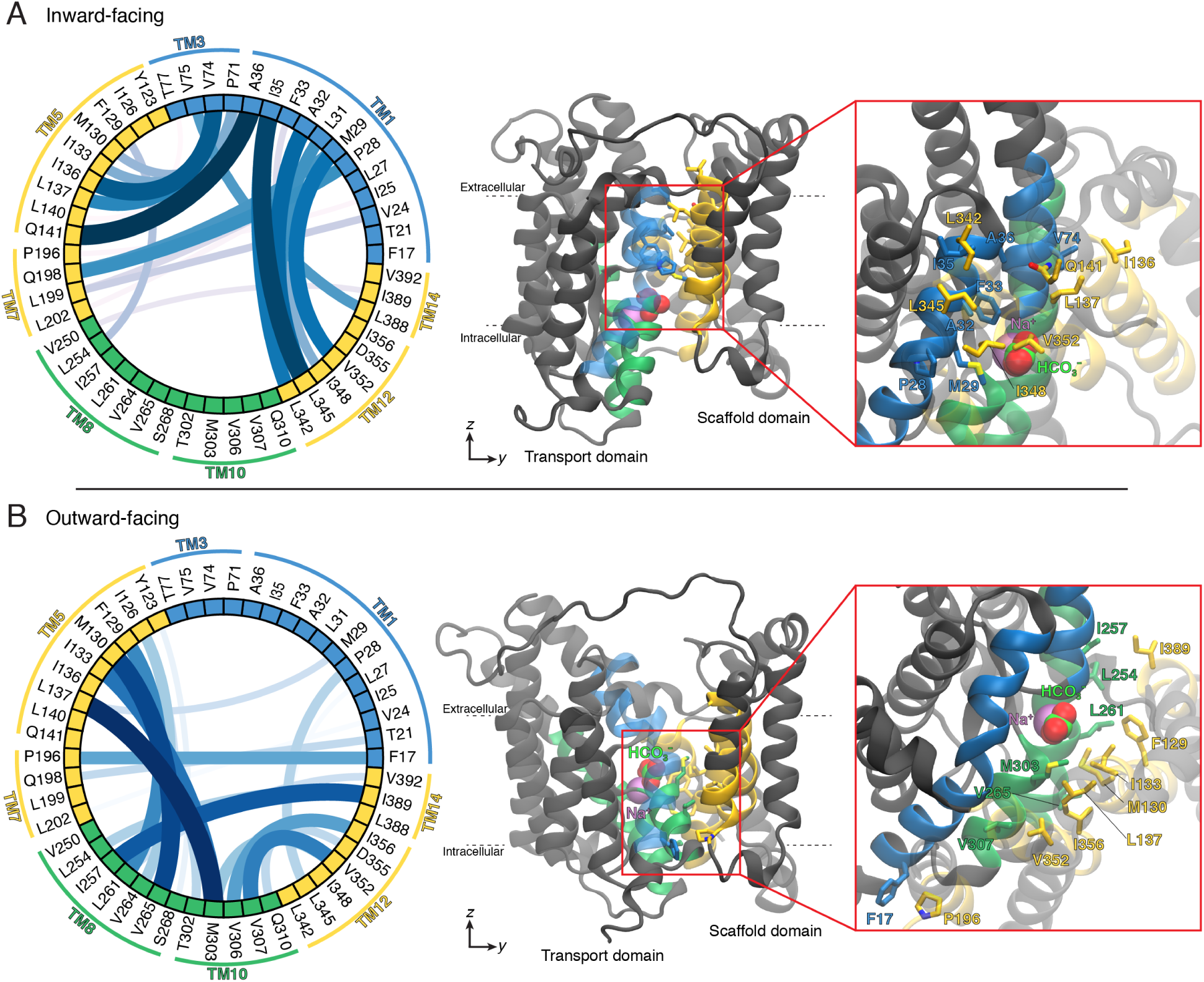
Hydrophobic gating residues of BicA. Chord diagram depicting the probability of interactions formed between gating residues in the (A) inward-facing conformation and (B) outward-facing conformation. Probability of interactions were calculated on 50,000 MD structures drawn from the respective free energy basin. Thickness and color intensity of connections between nodes represent the relative probability between two residues interacting. Accompanying MD snapshot showing selected residues mediating the closure of the transporter. STAS domain not shown for clarity. Transmembrane (TM) helices are colored as follows, TM1, TM3: blue; TM5, TM7, TM12, TM14: yellow; TM8, TM10: green.

Given that the molecular gates of BicA are primarily facilitated by the hydrophobic interactions of aliphatic side chains, we sought to determine if mutations may be introduced to increase the substrate transport rate. Specifically, we hypothesized if substitutions to alanine residues may decrease the contacting surface area, while still retaining the nonpolar local environment to maintain proper transport function. To this end, we targeted residues with large aliphatic side chains (Met, Leu, Ile, etc) located on various gating helices of BicA and residues of high contact probability based on the MD simulations. Three BicA triple mutants were simulated via the umbrella sampling protocol to delineate its effects on the energetics of the transporter conformational dynamics.

We first characterized mutations of residues on the hydrophobic gate that reside on the transport domain. The BicA triple mutant containing the substitutions Met29Ala;Phe33Ala;Ile35Ala are located on the extracellular half of transmembrane helix 1 (Figure 7, top). These residues were primarily found to interact with residues on TM12 to restrict access from the extracellular space and stabilize the inward-facing conformation (Figure 6A). The PMF profile of the Met29Ala;Phe33Ala;Ile35Ala BicA mutant reveals the free energy barrier to form the outward-facing state has decreased to 2.78 *±* 0.20 kcal/mol, as compared to the 3.97 *±* 0.24 kcal/mol in wild-type BicA (Figure 7, top). Furthermore, the relative free energy barrier of outward-facing to inward-facing transitions remains similar to the wild-type (Met29Ala;Phe33Ala;Ile35Ala: 0.56 kcal/mol, wildtype: 0.58 kcal/mol) consistent that these residues are primarily responsible for extracellular closure and not predicted to directly affect the stability of the outward-facing conformation.

**Figure 7:**
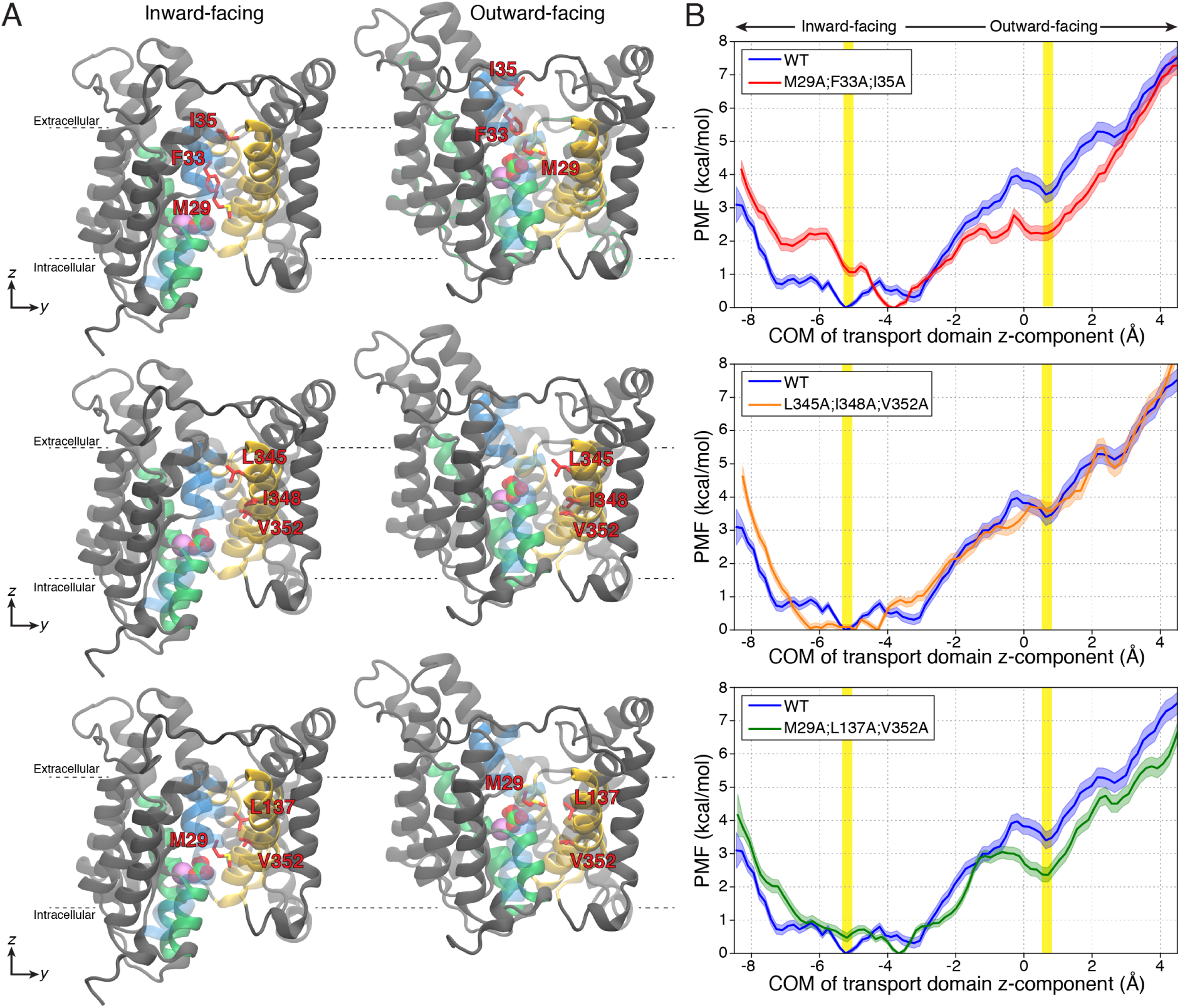
Simulated BicA triple mutants. (A) MD snapshots of the inward-facing and outward-facing conformations of BicA. Mutated residues are shown as sticks and colored red. The bicarbonate and sodium ion are shown as spheres and colored green and purple, respectively. Transmembrane (TM) helices are colored as follows, TM1, TM3: blue; TM5, TM7, TM12, TM14: yellow; TM8, TM10: green. STAS domain not shown for clarity. The mutants investigated in this study are Met29Ala;Phe33Ala;Ile35Ala (top), Leu345Ala;Ile348Ala;Val352Ala (middle), and Met29Ala;Leu137Ala,Val352Ala (bottom). (B) Potential of mean force (PMF) profiles of the three studied BicA triple mutants. The wild-type BicA is shown in blue and duplicated for comparison. BicA systems are colored as follows, wild-type: blue, Met29Ala;Phe33Ala;Ile35Ala: red, Leu345Ala;Ile348Ala;Val352Ala: orange, Met29Ala;Leu137Ala,Val352Ala: green. *x*-axis represents the *z*-coordinate displacement of the center of mass (COM) of the transport domain with respect to the center of mass of the scaffold domain. Inward- and outward-facing conformations are highlight as vertical yellow bars.

The residues of the second BicA triple mutant, Leu345Ala;Ile348Ala;Val352Ala, are located on transmembrane helix 12 of the scaffold domain (Figure 7, middle). The PMF profiles for Leu345Ala;Ile348Ala;Val352Ala predict a modest decrease in the free energy barrier (Leu345Ala;Ile348Ala;Val352Ala: 3.72 *±* 0.24 kcal/mol, wild-type: 3.97 *±* 0.24 kcal/mol). We suspect transmembrane helix 12 serves as a shared gating helix that facilitate both inward-facing and outward-facing conformations and contributes to a slight favorable reduction in the free energy barrier. However, these mutations also compromises on the destabilized interactions that close the intracellular pathway.

Lastly, the BicA mutant Met29Ala;Leu137Ala,Val352Ala targets residues on both the transport and scaffold domain (Figure 7, bottom). This mutant was specifically simulated to remove the hydrophobic interactions involving Met29. In the inward-facing state, Met29 interacts with Val352, whereas in the outward-facing state Met29 switches its interaction partner to Leu137 (Figure 6). Similar to the first described BicA triple mutant (Met29Ala;Phe33Ala;Ile35Ala), the PMF profile delineates a decrease in the free energy barrier (Met29Ala;Leu137Ala,Val352Ala: 3.05 *±* 0.19 kcal/mol, wild-type: 3.97 *±* 0.24 kcal/mol) and similar stability of the outward-facing state. In a multiple sequence alignment of 300 homologs generated by ConSurf, we find that the mutated residues are highly conserved in their physicochemical properties. The hydrophobic side chains of Met29, Ile35, and Ile348 are conserved, while Phe33 and Leu345 are entirely conserved. The MSA reveals that a population of BicA homologs have Thr in place of Val352 (Figure S23). Overall, our simulations predict alanine substitutions to residues on the transmembrane domain, specifically transmembrane helix 1, decrease the free energy barriers for structural rearrangements and expected to increase bicarbonate uptake.

## Conclusion

Bicarbonate transporters are key membrane transporters that regulate photosynthesis production. In this study, we utilized adaptive sampling and Markov state modeling to characterize the structural dynamics and thermodynamics of the BicA transport mechanism. Our simulations reveal that BicA protomers are more favored to undergo conformational transitions independently rather than simultaneously, consistent with previous cross-linking studies of the SLC26Dg transporter.^17^ In our simulations, we observed the cytoplasmic STAS domain does not remain stable in solution and undergo various structural rearrangements. We hypothesized, the stability of the STAS domain may be facilitated with other association proteins *in vivo*. Previous experimental characterization of the related *E. coli* YchM transporter has suggested the STAS domain to interact with a number of regulatory proteins with implications on transport activity.^77^ Further studies may focus on *in vivo* regulatory mechanisms of BicA.

We further investigated various BicA mutants that are predicted to affect the conformational energetics of the transport process. We predict that the unique steric constraints on the protein backbone provided by prolines residues located at junction of the scaffold and transport domain facilitate the proper conformational dynamics for transport. Furthermore, substitutions to bulky aliphatic residues that form the hydrophobic gate decrease the free energy barriers for inward-facing to outward-facing transitions. Specifically, alanine mutations located on transmembrane helix 1 are predicted to enhance transport with minimal consequences on the stability of conformations. Such mutations to the hydrophobic gating residues introduced to the sodium/proton antiporter PaNhaP were also identified to increase transport activity.^75^ However, how these mutations may affect transporter expression, folding, trafficking, or biogenesis cannot be delineated from simulations. In all, the extensive simulations conducted in this study provide a comprehensive mechanistic view of BicA transport dynamics and elucidate potential engineered mutations to enhance cyanobacteria photosynthetic yield.

## Author Contributions

M.C.C. and D.S. designed the study. D.S. acquired funding. M.C.C. and Y.A. performed simulations. M.C.C., Y.A., A.P., and D.S. analyzed data. D.S. supervised the study. M.C.C. and A.P. wrote manuscript with input from D.S. All authors reviewed the final manuscript.

## Acknowledgments

This research was part of the Blue Waters sustained-petascale computing project, which was supported by the National Science Foundation (awards OCI-0725070 and ACI-1238993) the State of Illinois, and as of December, 2019, the National Geospatial-Intelligence Agency. Blue Waters was a joint effort of the University of Illinois Urbana-Champaign and its National Center for Supercomputing Applications. D.S. acknowledges funding from the NSF CAREER Award MCB 18-45606 and funding from the National Institutes of Health (Award No. R35GM142745).

## Data Availability

Predicted outward-facing state of BicA, lipid parameters, and structures extracted from the Markov state model are available on the Zenodo repository: 10.5281/zenodo.6647550

## References

(1) Liu, H.; Nolla, H. A.; Campbell, L. Prochlorococcus growth rate and contribution to primary production in the equatorial and subtropical North Pacific Ocean. Aquatic Microbial Ecology 1997, 12, 39–47.

(2) Partensky, F.; Hess, W. R.; Vaulot, D. Prochlorococcus, a marine photosynthetic prokaryote of global significance. Microbiology and molecular biology reviews 1999, 63, 106–127.

(3) Price, G. D.; Woodger, F. J.; Badger, M. R.; Howitt, S. M.; Tucker, L. Identification of a SulP-type bicarbonate transporter in marine cyanobacteria. Proceedings of the National Academy of Sciences 2004, 101, 18228–18233.

(4) Price, G. D. Inorganic carbon transporters of the cyanobacterial CO 2 concentrating mechanism. Photosynthesis research 2011, 109, 47–57.

(5) Wang, C.; Sun, B.; Zhang, X.; Huang, X.; Zhang, M.; Guo, H.; Chen, X.; Huang, F.; Chen, T.; Mi, H.; Yu, F.; Liu, L.-N.; Zhang, P. Structural mechanism of the active bicarbonate transporter from cyanobacteria. Nature Plants 2019, 5, 1184–1193.

(6) Liu, X.-Y.; Hou, W.-T.; Wang, L.; Li, B.; Chen, Y.; Chen, Y.; Jiang, Y.-L.; Zhou, C.-Z. Structures of cyanobacterial bicarbonate transporter SbtA and its complex with PII-like SbtB. Cell Discovery 2021, 7, 1–5.

(7) Bracher, A.; Whitney, S. M.; Hartl, F. U.; Hayer-Hartl, M. Biogenesis and metabolic maintenance of Rubisco. Annual review of plant biology 2017, 68, 29–60.

(8) Rae, B. D.; Long, B. M.; Förster, B.; Nguyen, N. D.; Velanis, C. N.; Atkinson, N.; Hee, W. Y.; Mukherjee, B.; Price, G. D.; McCormick, A. J. Progress and challenges of engineering a biophysical CO2-concentrating mechanism into higher plants. Journal of Experimental Botany 2017, 68, 3717–3737.

(9) Badger, M. R.; Price, G. D. CO2 concentrating mechanisms in cyanobacteria: molecular components, their diversity and evolution. Journal of experimental botany 2003, 54, 609–622.

(10) McGrath, J. M.; Long, S. P. Can the cyanobacterial carbon-concentrating mechanism increase photosynthesis in crop species? A theoretical analysis. Plant physiology 2014, 164, 2247–2261.

(11) Du, J.; Förster, B.; Rourke, L.; Howitt, S. M.; Price, G. D. Characterisation of cyanobacterial bicarbonate transporters in E. coli shows that SbtA homologs are functional in this heterologous expression system. PloS one 2014, 9, e115905.

(12) Rolland, V.; Badger, M. R.; Price, G. D. Redirecting the cyanobacterial bicarbonate transporters BicA and SbtA to the chloroplast envelope: soluble and membrane cargos need different chloroplast targeting signals in plants. Frontiers in plant science 2016, 7, 185.

(13) Gupta, J. K.; Rai, P.; Jain, K. K.; Srivastava, S. Overexpression of bicarbonate transporters in the marine cyanobacterium Synechococcus sp. PCC 7002 increases growth rate and glycogen accumulation. Biotechnology for biofuels 2020, 13, 1–12.

(14) Chang, Y.-N.; Geertsma, E. R. The novel class of seven transmembrane segment inverted repeat carriers. Biological Chemistry 2017, 398, 165–174.

(15) Gorbunov, D.; Sturlese, M.; Nies, F.; Kluge, M.; Bellanda, M.; Battistutta, R.; Oliver, D. Molecular architecture and the structural basis for anion interaction in prestin and SLC26 transporters. Nature communications 2014, 5, 3622.

(16) Walter, J. D.; Sawicka, M.; Dutzler, R. Cryo-EM structures and functional characterization of murine Slc26a9 reveal mechanism of uncoupled chloride transport. Elife 2019, 8, e46986.

(17) Chang, Y.-N.; Jaumann, E. A.; Reichel, K.; Hartmann, J.; Oliver, D.; Hummer, G.; Joseph, B.; Geertsma, E. R. Structural basis for functional interactions in dimers of SLC26 transporters. Nature Communications 2019, 10, 2032.

(18) Yu, X.; Yang, G.; Yan, C.; Baylon, J. L.; Jiang, J.; Fan, H.; Lu, G.; Hasegawa, K.; Okumura, H.; Wang, T.; Tajkhorshid, E.; Li, S.; Yan, N. Dimeric structure of the uracil:proton symporter UraA provides mechanistic insights into the SLC4/23/26 transporters. Cell Research 2017, 27, 1020–1033.

(19) Wang, W.; Tsirulnikov, K.; Zhekova, H. R.; Kayık, G.; Khan, H. M.; Azimov, R.; Abuladze, N.; Kao, L.; Newman, D.; Noskov, S. Y.; Zhou, Z. H.; Pushkin, A.; Kurtz, I. Cryo-EM structure of the sodium-driven chloride/bicarbonate exchanger NDCBE. Nature Communications 2021, 12, 5690.

(20) Coudray, N.; L. Seyler, S.; Lasala, R.; Zhang, Z.; Clark, K. M.; Dumont, M. E.; Rohou, A.; Beckstein, O.; Stokes, D. L. Structure of the SLC4 transporter Bor1p in an inward-facing conformation. Protein Science 2017, 26, 130–145.

(21) Jardetzky, O. Simple allosteric model for membrane pumps. Nature 1966, 211, 969–970.

(22) Alper, S. L.; Sharma, A. K. The SLC26 gene family of anion transporters and channels. Molecular aspects of medicine 2013, 34, 494–515.

(23) Long, S. P.; Marshall-Colon, A.; Zhu, X.-G. Meeting the global food demand of the future by engineering crop photosynthesis and yield potential. Cell 2015, 161, 56–66.

(24) Searchinger, T.; Waite, R.; Hanson, C.; Ranganathan, J.; Matthews, E. Creating a Sustainable Food Future: A Menu of Solutions to Feed Nearly 10 Billion People by 2050; 2019.

(25) Hollingsworth, S. A.; Dror, R. O. Molecular Dynamics Simulation for All. Neuron 2018, 99, 1129–1143.

(26) Chan, M. C.; Shukla, D. Markov state modeling of membrane transport proteins. Journal of Structural Biology 2021, 213, 107800.

(27) Chen, J.; White, A.; Nelson, D. C.; Shukla, D. Role of substrate recognition in modulating strigolactone receptor selectivity in witchweed. Journal of Biological Chemistry 2021, 297, 101092.

(28) Shukla, S.; Zhao, C.; Shukla, D. Dewetting Controls Plant Hormone Perception and Initiation of Drought Resistance Signaling. Structure 2019, 27, 692–702.e3.

(29) Moffett, A. S.; Bender, K. W.; Huber, S. C.; Shukla, D. Molecular dynamics simulations reveal the conformational dynamics of Arabidopsis thaliana BRI1 and BAK1 receptorlike kinases. Journal of Biological Chemistry 2017, 292, 12643–12652.

(30) Zhou, H.; Dong, Z.; Verkhivker, G.; Zoltowski, B. D.; Tao, P. Allosteric mechanism of the circadian protein Vivid resolved through Markov state model and machine learning analysis. PLOS Computational Biology 2019, 15, e1006801.

(31) Zhao, Y.; Zhang, Y.; Sun, M.; Zheng, Q. A theoretical study on the signal transduction process of bacterial photoreceptor PpSB1 based on the Markov state model. Physical Chemistry Chemical Physics 2021, 23, 2398–2405.

(32) Trozzi, F.; Wang, F.; Verkhivker, G.; Zoltowski, B. D.; Tao, P. Dimeric allostery mechanism of the plant circadian clock photoreceptor ZEITLUPE. PLOS Computational Biology 2021, 17, e1009168.

(33) Selvam, B.; Yu, Y.-C.; Chen, L.-Q.; Shukla, D. Molecular Basis of the Glucose Transport Mechanism in Plants. ACS Central Science 2019, 5, 1085–1096.

(34) Cheng, K. J.; Selvam, B.; Chen, L.-Q.; Shukla, D. Distinct Substrate Transport Mechanism Identified in Homologous Sugar Transporters. The Journal of Physical Chemistry B 2019, 123, 8411–8418.

(35) Weigle, A. T.; Carr, M.; Shukla, D. Impact of Increased Membrane Realism on Conformational Sampling of Proteins. Journal of Chemical Theory and Computation 2021, 17, 5342–5357.

(36) Feng, J.; Selvam, B.; Shukla, D. How do antiporters exchange substrates across the cell membrane? An atomic-level description of the complete exchange cycle in NarK. Structure 2021, 29, 922–933.e3.

(37) Chan, M. C.; Procko, E.; Shukla, D. Structural Rearrangement of the Serotonin Transporter Intracellular Gate Induced by Thr276 Phosphorylation. ACS Chemical Neuroscience 2022, 13, 933–945.

(38) Selvam, B.; Mittal, S.; Shukla, D. Free Energy Landscape of the Complete Transport Cycle in a Key Bacterial Transporter. ACS Central Science 2018, 4, 1146–1154.

(39) Beckstein, O.; Naughton, F. General principles of secondary active transporter function. Biophysics Reviews 2022, 3, 011307.

(40) Drew, D.; Boudker, O. Shared Molecular Mechanisms of Membrane Transporters. Annual Review of Biochemistry 2016, 85, 543–572.

(41) Matsuoka, R.; Fudim, R.; Jung, S.; Zhang, C.; Bazzone, A.; Chatzikyriakidou, Y.; Robinson, C. V.; Nomura, N.; Iwata, S.; Landreh, M.; Orellana, L.; Beckstein, O.; Drew, D. Structure, mechanism and lipid-mediated remodeling of the mammalian Na+/H+ exchanger NHA2. Nature Structural & Molecular Biology 2022, 29, 108–120.

(42) Winkelmann, I.; Matsuoka, R.; Meier, P. F.; Shutin, D.; Zhang, C.; Orellana, L.; Sexton, R.; Landreh, M.; Robinson, C. V.; Beckstein, O.; Drew, D. Structure and elevator mechanism of the mammalian sodium/proton exchanger NHE9. The EMBO Journal 2020, 39, 4541–4559.

(43) Lee, C.; Kang, H. J.; von Ballmoos, C.; Newstead, S.; Uzdavinys, P.; Dotson, D. L.; Iwata, S.; Beckstein, O.; Cameron, A. D.; Drew, D. A two-domain elevator mechanism for sodium/proton antiport. Nature 2013, 501, 573–577.

(44) Coincon, M.; Uzdavinys, P.; Nji, E.; Dotson, D. L.; Winkelmann, I.; Abdul-Hussein, S.; Cameron, A. D.; Beckstein, O.; Drew, D. Crystal structures reveal the molecular basis of ion translocation in sodium/proton antiporters. Nature Structural & Molecular Biology 2016, 23, 248–255.

(45) Dong, Y.; Gao, Y.; Ilie, A.; Kim, D.; Boucher, A.; Li, B.; Zhang, X. C.; Orlowski, J.; Zhao, Y. Structure and mechanism of the human NHE1-CHP1 complex. Nature Communications 2021, 12, 3474.

(46) Ruan, Y.; Miyagi, A.; Wang, X.; Chami, M.; Boudker, O.; Scheuring, S. Direct visualization of glutamate transporter elevator mechanism by high-speed AFM. Proceedings of the National Academy of Sciences 2017, 114, 1584–1588.

(47) Martínez, L.; Andrade, R.; Birgin, E. G.; Martínez, J. M. PACKMOL: A package for building initial configurations for molecular dynamics simulations. Journal of Computational Chemistry 2009, 30, 2157–2164.

(48) Gomez, Y. K.; Natale, A. M.; Lincoff, J.; Wolgemuth, C. W.; Rosenberg, J. M.; Grabe, M. Taking the Monte-Carlo gamble: How not to buckle under the pressure! Journal of Computational Chemistry 2021, 43, 431–434.

(49) Webb, B.; Sali, A. Comparative Protein Structure Modeling Using MODELLER. Current Protocols in Bioinformatics 2016, 54, 5.6.1–5.6.37.

(50) Jumper, J. et al. Highly accurate protein structure prediction with AlphaFold. Nature 2021, 596, 583–589.

(51) Evans, R. et al. Protein complex prediction with AlphaFold-Multimer. bioRxiv 2021, doi:10.1101/2021.10.04.463034.

(52) Hopkins, C. W.; Le Grand, S.; Walker, R. C.; Roitberg, A. E. Long-time-step molecular dynamics through hydrogen mass repartitioning. Journal of chemical theory and computation 2015, 11, 1864–1874.

(53) Phillips, J. C. et al. Scalable molecular dynamics on CPU and GPU architectures with NAMD. The Journal of Chemical Physics 2020, 153, 044130.

(54) Huynh, K. W.; Jiang, J.; Abuladze, N.; Tsirulnikov, K.; Kao, L.; Shao, X.; Newman, D.; Azimov, R.; Pushkin, A.; Zhou, Z. H.; Kurtz, I. CryoEM structure of the human SLC4A4 sodium-coupled acid-base transporter NBCe1. Nature Communications 2018, 9, 900.

(55) Roe, D. R.; Cheatham, T. E. PTRAJ and CPPTRAJ: Software for Processing and Analysis of Molecular Dynamics Trajectory Data. J. Chem. Theory Comput. 2013, 9, 3084–3095.

(56) McGibbon, R.; Beauchamp, K.; Harrigan, M.; Klein, C.; Swails, J.; Herńandez, C.; Schwantes, C.; Wang, L.-P.; Lane, T.; Pande, V. MDTraj: A Modern Open Library for the Analysis of Molecular Dynamics Trajectories. Biophys. J 2015, 109, 1528–1532.

(57) Humphrey, W.; Dalke, A.; Schulten, K. VMD: visual molecular dynamics. Journal of molecular graphics 1996, 14, 33–38.

(58) Grant, B. J.; Rodrigues, A. P.; ElSawy, K. M.; McCammon, J. A.; Caves, L. S. Bio3d: an R package for the comparative analysis of protein structures. Bioinformatics 2006, 22, 2695–2696.

(59) Scherer, M. K.; Trendelkamp-Schroer, B.; Paul, F.; Pérez-Herández, G.; Hoffmann, M.; Plattner, N.; Wehmeyer, C.; Prinz, J.-H.; Nóe, F. PyEMMA 2: A Software Package for Estimation, Validation, and Analysis of Markov Models. Journal of Chemical Theory and Computation 2015, 11, 5525–5542.

(60) Zhou, Q. et al. Common activation mechanism of class A GPCRs. eLife 2019, 8, e50279.

(61) Shirts, M. R.; Chodera, J. D. Statistically optimal analysis of samples from multiple equilibrium states. The Journal of chemical physics 2008, 129, 124105.

(62) Ashkenazy, H.; Abadi, S.; Martz, E.; Chay, O.; Mayrose, I.; Pupko, T.; Ben-Tal, N. ConSurf 2016: an improved methodology to estimate and visualize evolutionary conservation in macromolecules. Nucleic acids research 2016, 44, W344–W350.

(63) Crooks, G. E.; Hon, G.; Chandonia, J.-M.; Brenner, S. E. WebLogo: a sequence logo generator. Genome research 2004, 14, 1188–1190.

(64) Murata, N.; Wada, H.; Gombos, Z. Modes of Fatty-Acid Desaturation in Cyanobacteria. Plant and Cell Physiology 1992, 33, 933–941.

(65) Wada, H.; Murata, N. Lipids in photosynthesis: structure, function and genetics; Springer, 1998; pp 65–81.

(66) Case, D. A., et al. AMBER 2018. University of California, San Francisco 2018

(67) Klauda, J. B.; Venable, R. M.; Freites, J. A.; O’Connor, J. W.; Tobias, D. J.; Mondragon-Ramirez, C.; Vorobyov, I.; MacKerell, A. D.; Pastor, R. W. Update of the CHARMM All-Atom Additive Force Field for Lipids: Validation on Six Lipid Types. The Journal of Physical Chemistry B 2010, 114, 7830–7843.

(68) van Eerden, F. J.; de Jong, D. H.; de Vries, A. H.; Wassenaar, T. A.; Marrink, S. J. Characterization of thylakoid lipid membranes from cyanobacteria and higher plants by molecular dynamics simulations. Biochimica et Biophysica Acta (BBA) - Biomembranes 2015, 1848, 1319–1330.

(69) Shibagaki, N.; Grossman, A. R. Binding of Cysteine Synthase to the STAS Domain of Sulfate Transporter and Its Regulatory Consequences. Journal of Biological Chemistry 2010, 285, 25094–25102.

(70) Ko, S. B.; Zeng, W.; Dorwart, M. R.; Luo, X.; Kim, K. H.; Millen, L.; Goto, H.; Naruse, S.; Soyombo, A.; Thomas, P. J.; Muallem, S. Gating of CFTR by the STAS domain of SLC26 transporters. Nature Cell Biology 2004, 6, 343–350.

(71) Jiang, T.; Wen, P.-C.; Trebesch, N.; Zhao, Z.; Pant, S.; Kapoor, K.; Shekhar, M.; Tajkhorshid, E. Computational Dissection of Membrane Transport at a Microscopic Level. Trends in Biochemical Sciences 2020, 45, 202–216.

(72) Dror, R. O.; Dirks, R. M.; Grossman, J.; Xu, H.; Shaw, D. E. Biomolecular Simulation: A Computational Microscope for Molecular Biology. Annual Review of Biophysics 2012, 41, 429–452.

(73) Husic, B. E.; Pande, V. S. Markov state models: From an art to a science. Journal of the American Chemical Society 2018, 140, 2386–2396.

(74) Masrati, G.; Mondal, R.; Rimon, A.; Kessel, A.; Padan, E.; Lindahl, E.; Ben-Tal, N. An angular motion of a conserved four-helix bundle facilitates alternating access transport in the TtNapA and EcNhaA transporters. Proceedings of the National Academy of Sciences 2020, 117, 31850–31860.

(75) Okazaki, K.-i.; Wöhlert, D.; Warnau, J.; Jung, H.; Yildiz, Ö.; Kühlbrandt, W.; Hummer, G. Mechanism of the electroneutral sodium/proton antiporter PaNhaP from transition-path shooting. Nature communications 2019, 10, 1742.

(76) Jiang, T.; Wen, P.-C.; Trebesch, N.; Zhao, Z.; Pant, S.; Kapoor, K.; Shekhar, M.; Tajkhorshid, E. Computational dissection of membrane transport at a microscopic level. Trends in biochemical sciences 2020, 45, 202–216.

(77) Babu, M.; Greenblatt, J. F.; Emili, A.; Strynadka, N. C.; Reithmeier, R. A.; Moraes, T. F. Structure of a SLC26 anion transporter STAS domain in complex with acyl carrier protein: implications for E. coli YchM in fatty acid metabolism. Structure 2010, 18, 1450–1462.

