## Supporting Information for "Elevator-type Mechanism of the Cyanobacterial Bicarbonate Transporter"

#### Supplemental Figures

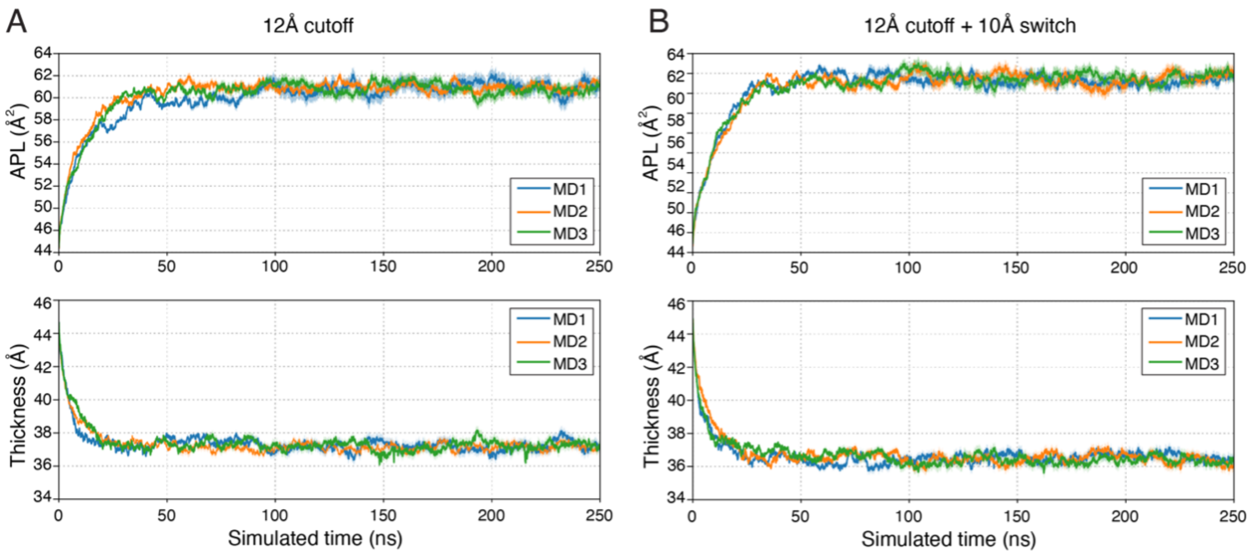


**Figure S1: Stability of the membrane in simulations.** Time-resolved measurements, area-per-lipid (APL) and membrane thickness of the cyanobacteria lipid membrane. Membrane stability using a 10Å potential-switching and Monte Carlo barostat. Three replicates of the cyanobacterial plasma membrane were simulated using either a (A) 12Å nonbonded cutoff or a (B) 12Å cutoff with a 10Å potential switch.

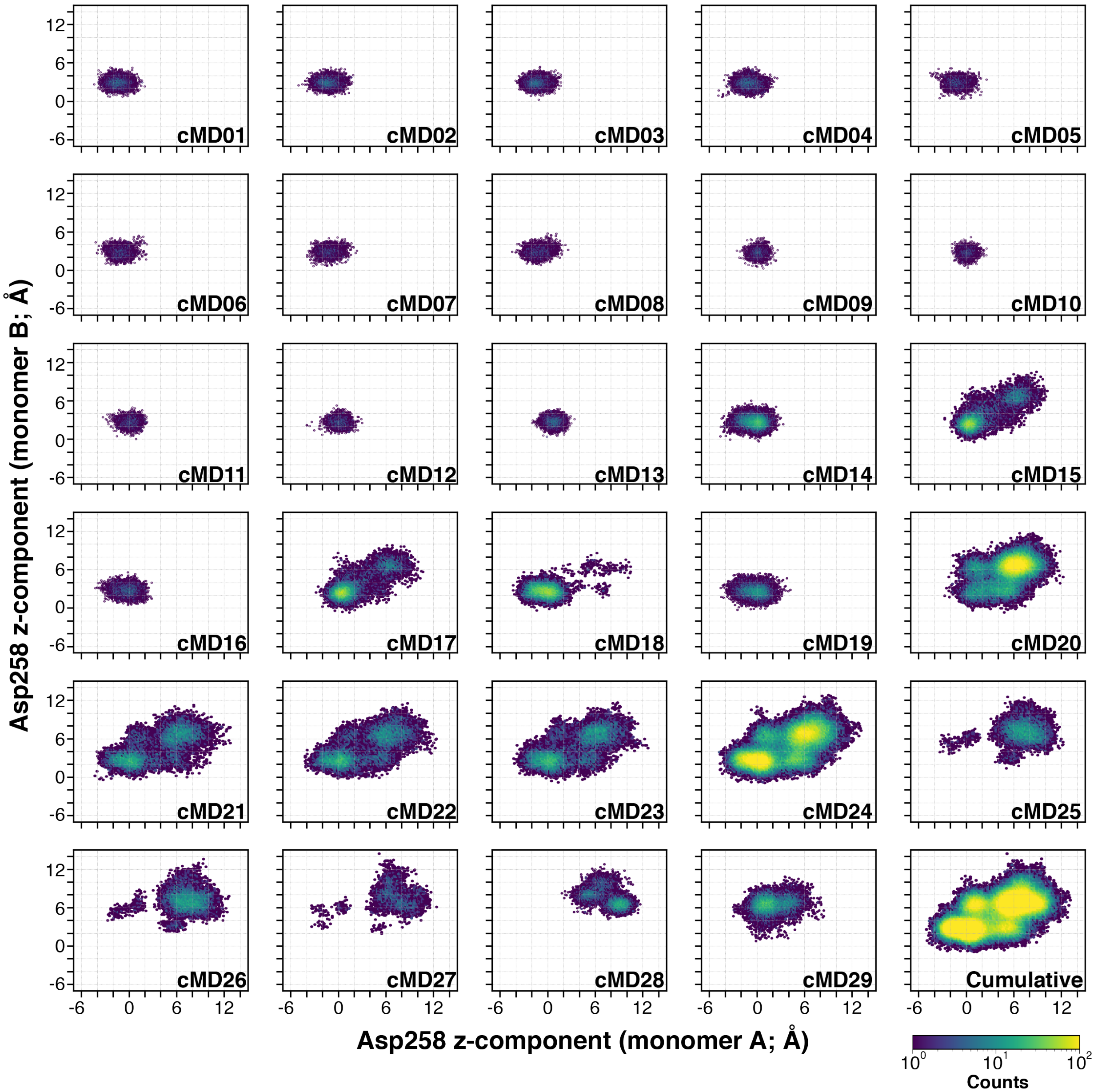


**Figure S2: Progression of the adaptive sampling simulation.** The $z$-coordinates of the C$\alpha$ atom of Asp258 from each adaptive round are projected as a two-dimensional histogram. The adaptive sampling round is indicated in the bottom right of each subplot.

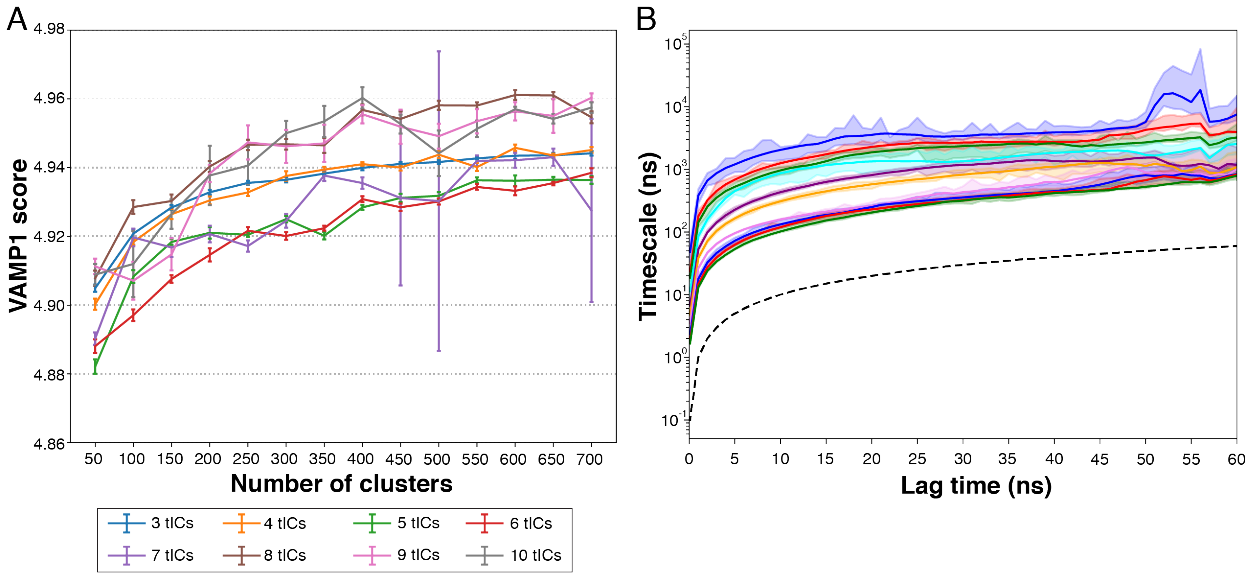


**Figure S3: Hyperparameter optimization for MSM construction**. (A) VAMP1 scores as a function of clustering size and tICA (time independent component analysis) decomposition. The highest scoring model was achieved with 400 clusters and 10 time-independent components. (B) Implied timescale plots from the transition probability matrix of the MSM. The first ten eigenvalues of the transition matrix which corresponds to the ten slowest transition rates is plotted for the optimized hyperparameters of the simulation datasets. The implied timescale curves converge at ∼10 ns lag time suggesting Markovian nature. Final models were constructed at lag time of 10 ns.

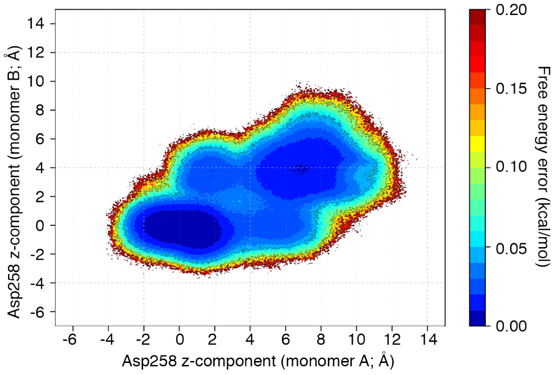


**Figure S4: Standard error measurements of the conformational free energy landscape**. The standard error of the free energy landscape was calculated using the bootstrap method by randomly selecting 80% of the trajectories for 500 independent samples and then projected on the coordinates of the C$\alpha$ atom of Asp258 for the two BicA protomers.

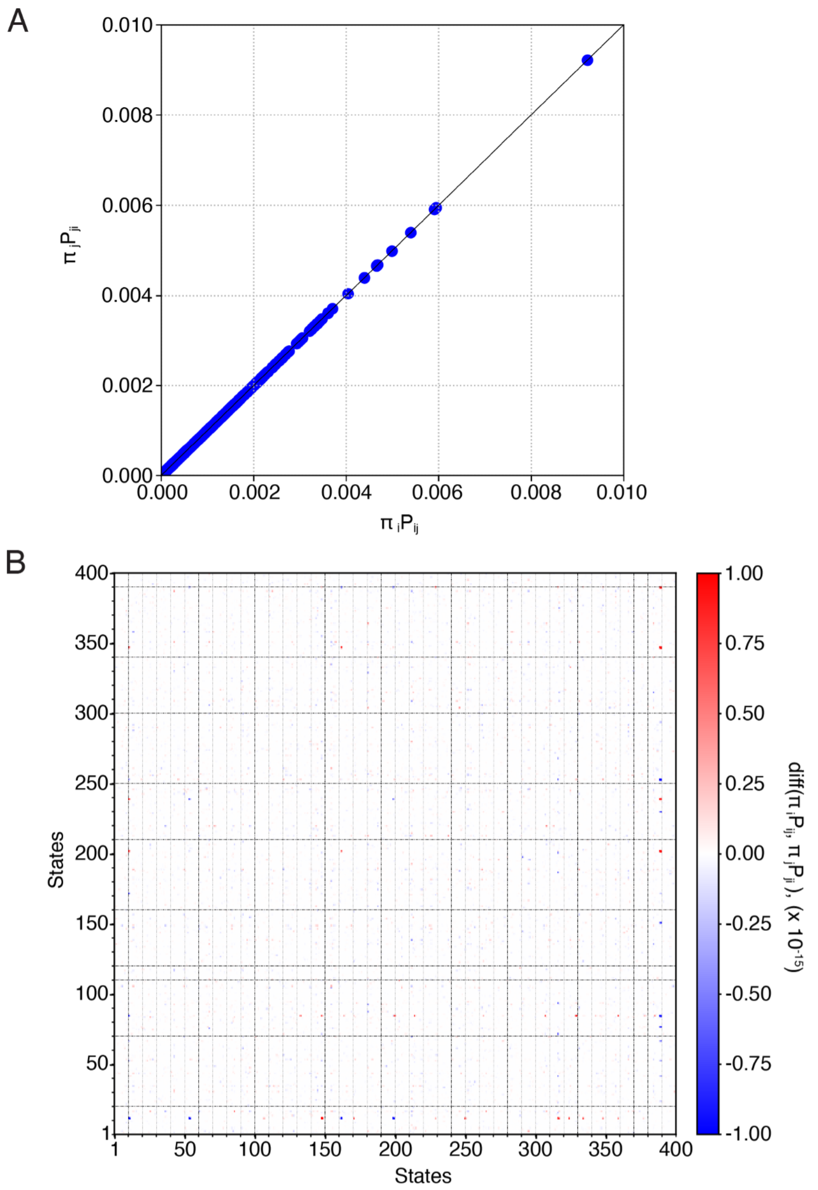


**Figure S5: Reversibility of the MSM transition probability matrix.** The reversibility was assessed if condition $\pi_i P_{ij} = \pi_j P_{ji}$ held for all MSM states i and j. (A) The quantities $\pi_i P_{ij}$ and $\pi_j P_{ji}$ plotted against each other for all MSM states. (B) The difference between $\pi_i P_{ij}$ and $\pi_j P_{ji}$ showing the two quantities vary within $10^{-15}$.

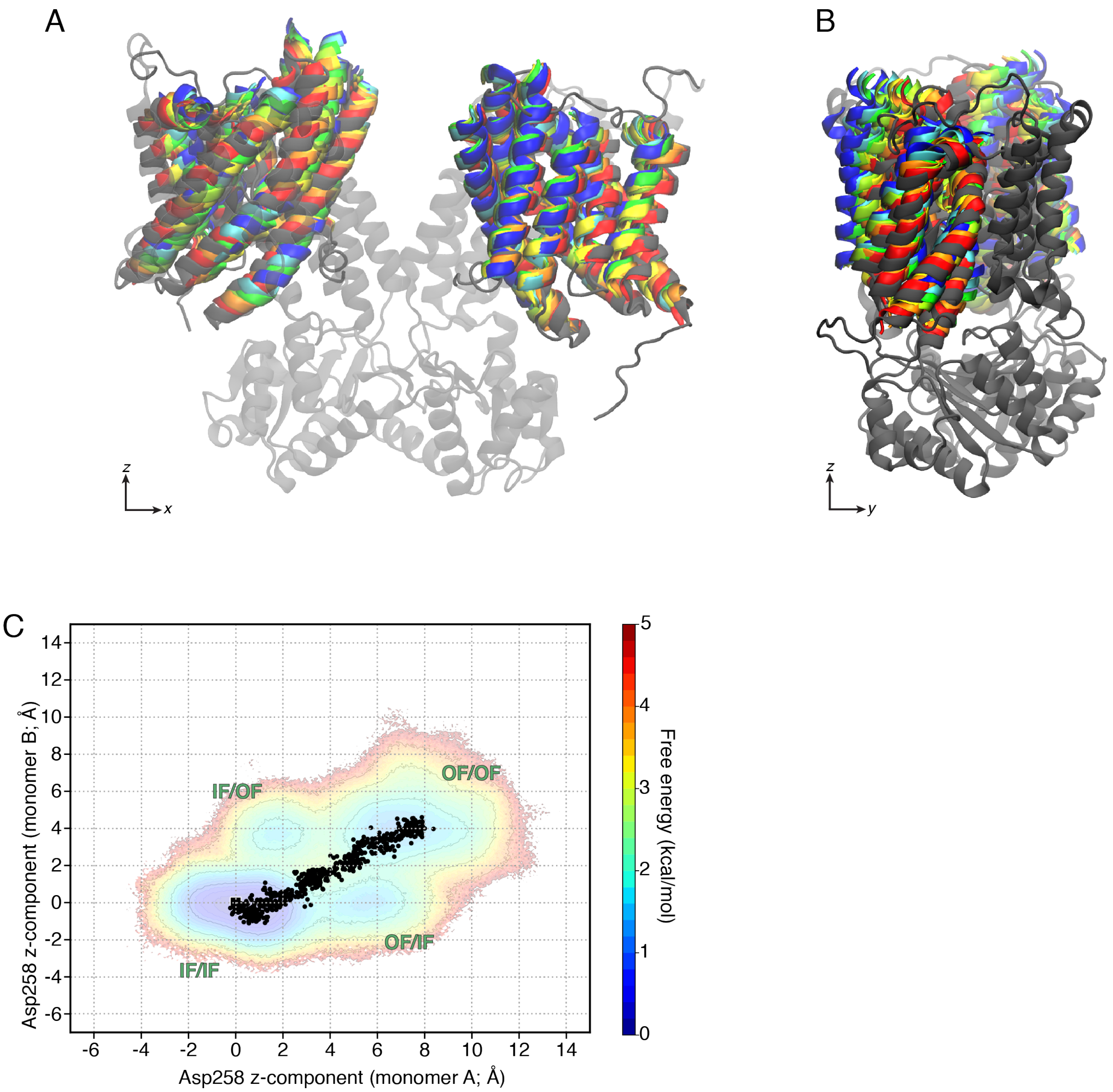


**Figure S6: Targeted MD trajectory to initiate sampling from the inward-facing to outward-facing state**. (A, B) Progression of the transport domain from targeted MD trajectory from in the inward-facing state (grey) to the outward-facing state (blue). The scaffold and STAS domain is colored in grey and rendered as transparent for clarity. (C) The progression of the $z$-coordinate of the C$\alpha$ atom of Asp258 during the targeted MD trajectory projected on the free energy landscape.

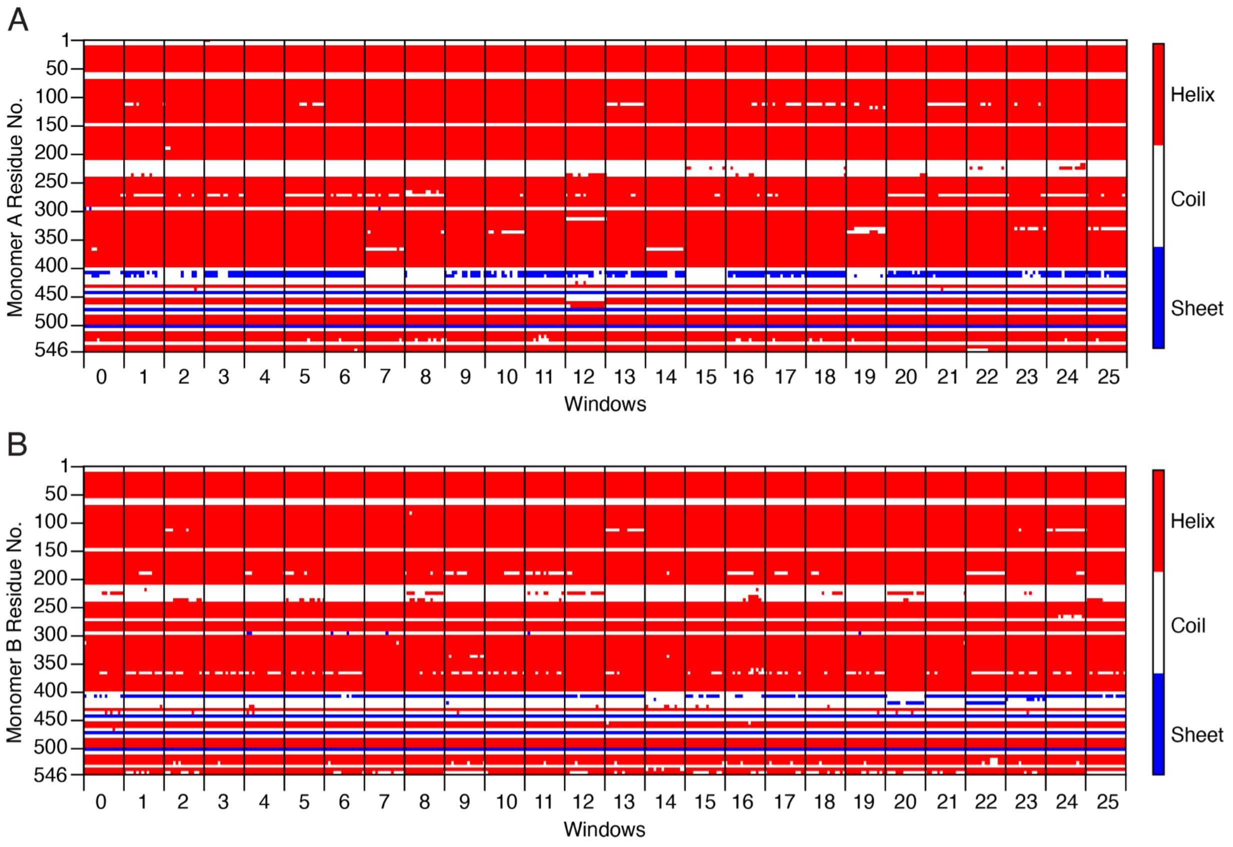


**Figure S7: DSSP secondary structure assignment of wild-type BicA residues across umbrella sampling windows. Each 16ns US simulation is plotted for each window. Secondary structures of each residue are assigned by color: red: helix, white: coil, and blue: sheet.**

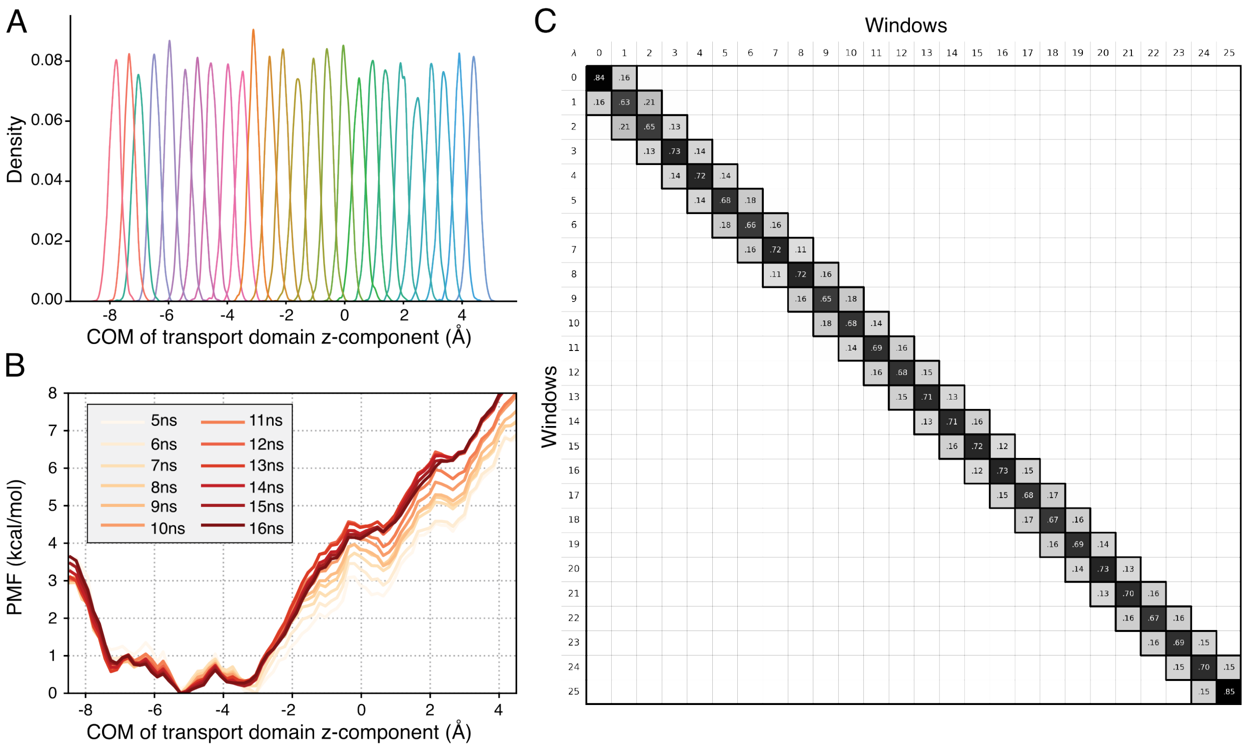


**Figure S8: Convergence of the wild-type BicA umbrella sampling simulations**. (A) Histogram of umbrella sampling windows along the reaction coordinate (B) Convergence of the PMF from 5ns to 16ns of simulation time. (C) Overlap matrix of umbrella sampling windows.

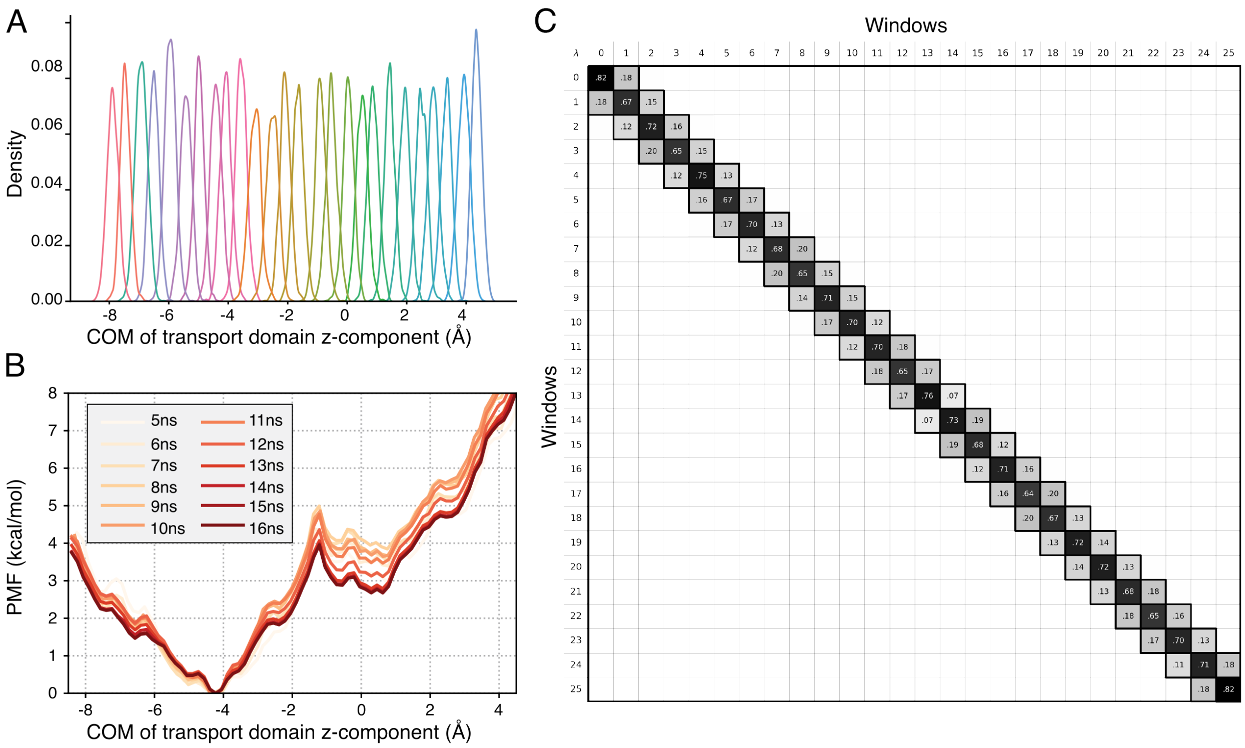


**Figure S9: Convergence of the BicA P341G-P122G mutant umbrella sampling simulations**. (A) Histogram of umbrella sampling windows along the reaction coordinate (B) Convergence of the PMF from 5ns to 16ns of simulation time. (C) Overlap matrix of umbrella sampling windows.

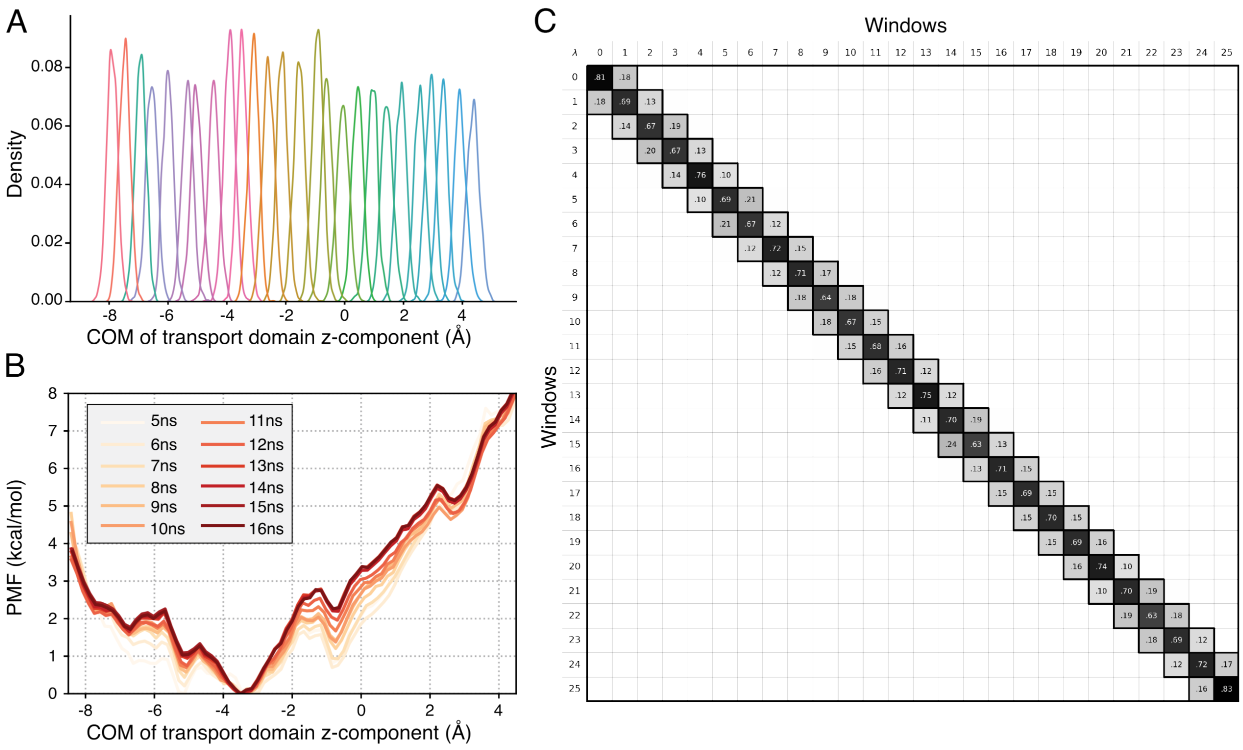


**Figure S10: Convergence of the BicA P341A-P122A mutant umbrella sampling simulations**. (A) Histogram of umbrella sampling windows along the reaction coordinate (B) Convergence of the PMF from 5ns to 16ns of simulation time. (C) Overlap matrix of umbrella sampling windows.

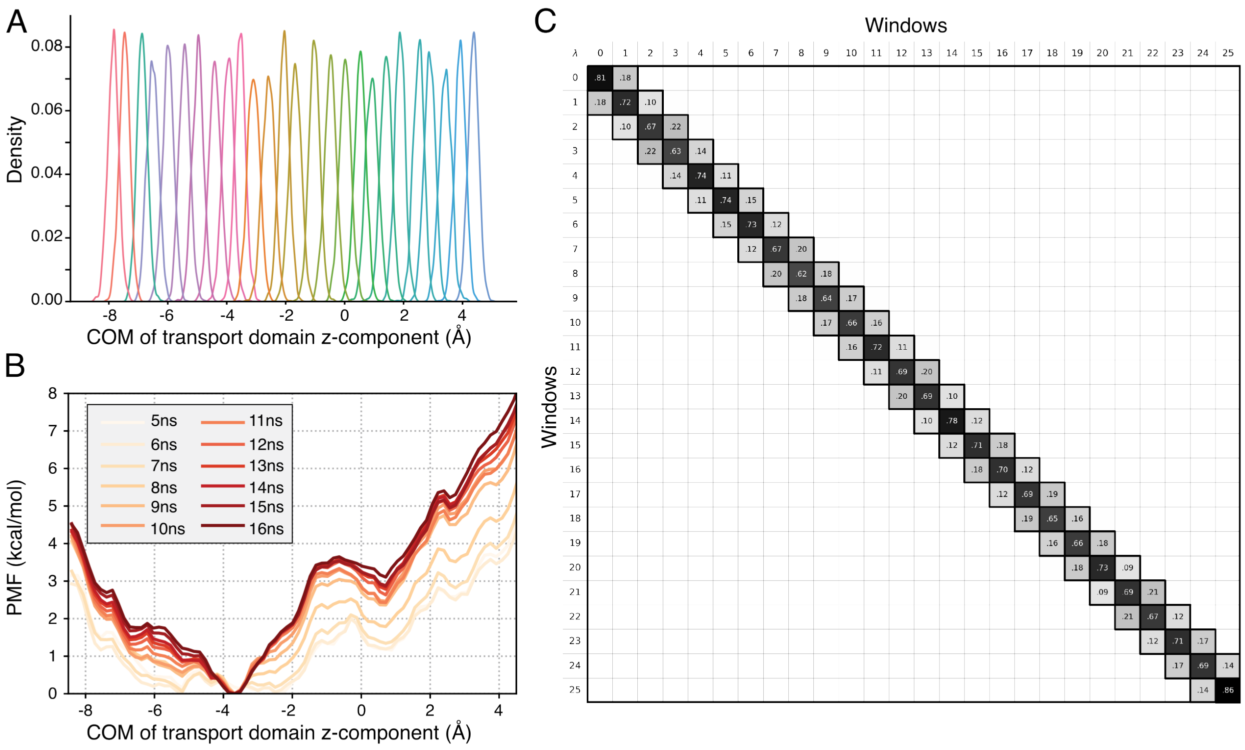


**Figure S11: Convergence of the BicA M29A-V352A-L137A mutant umbrella sampling simulations**. (A) Histogram of umbrella sampling windows along the reaction coordinate (B) Convergence of the PMF from 5ns to 16ns of simulation time. (C) Overlap matrix of umbrella sampling windows.

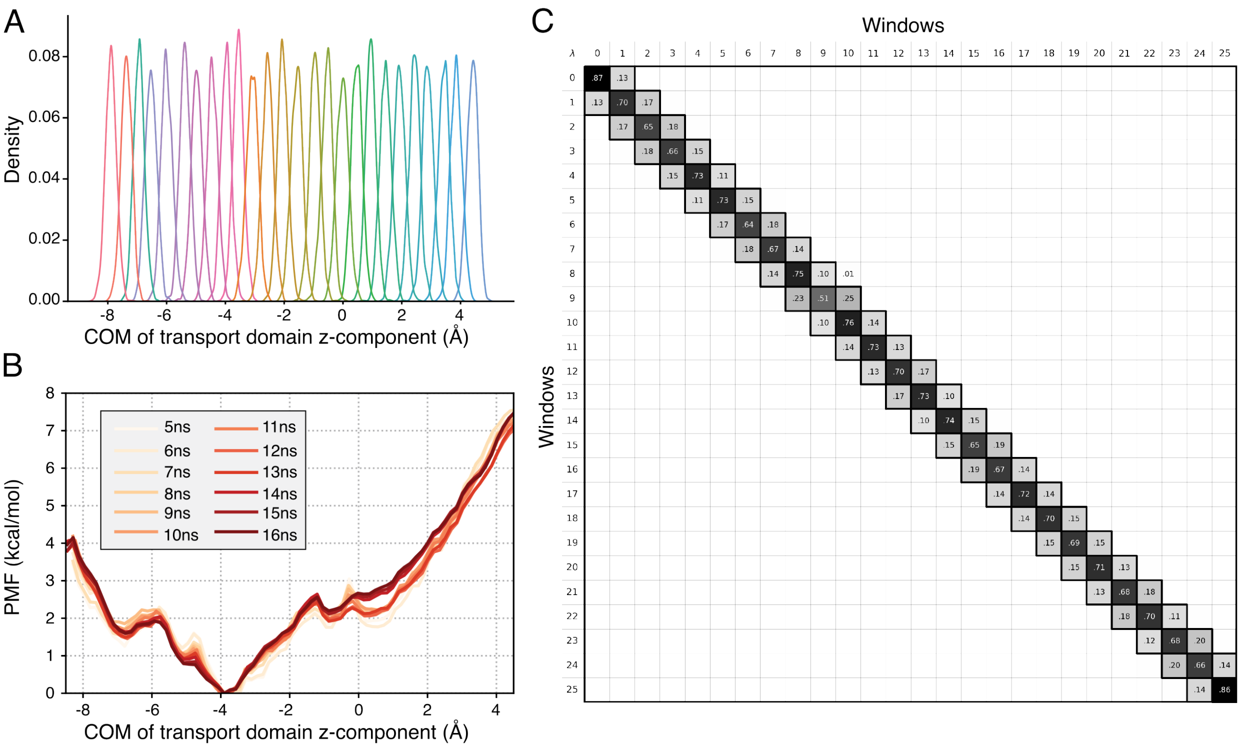


**Figure S12: Convergence of the BicA M29A-F33A-I35A mutant umbrella sampling simulations**. (A) Histogram of umbrella sampling windows along the reaction coordinate (B) Convergence of the PMF from 5ns to 16ns of simulation time. (C) Overlap matrix of umbrella sampling windows.

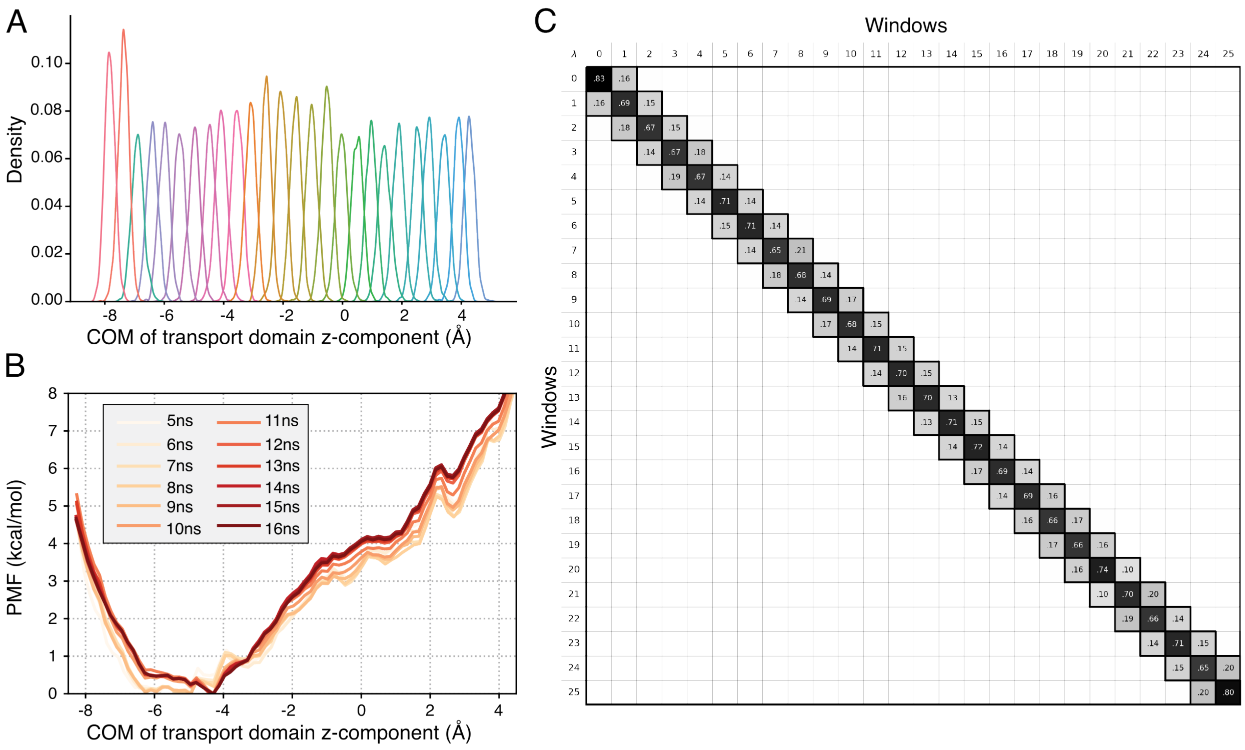


**Figure S13: Convergence of the BicA L345A-I348A-V352A mutant umbrella sampling simulations**. (A) Histogram of umbrella sampling windows along the reaction coordinate (B) Convergence of the PMF from 5ns to 16ns of simulation time. (C) Overlap matrix of umbrella sampling windows.

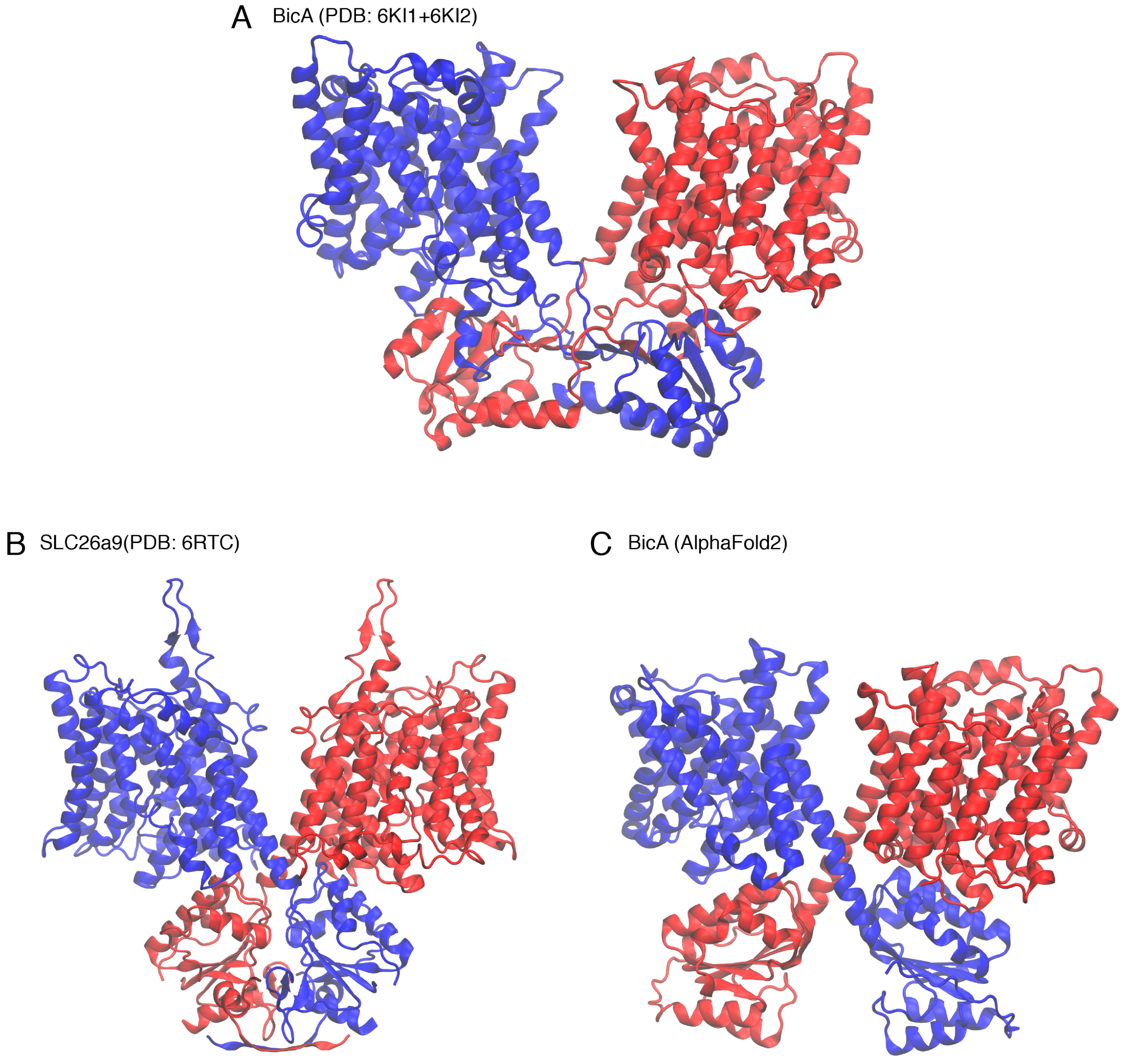


**Figure S14: Full-length models of the BicA dimer.** (A) BicA dimer model constructing using the two structural structures PDB: 6KI1 and PDB: 6KI2. The two monomers are colored red and blue, respectively. (B) The resolved related SLC26a9 transporter (PDB: 6RTC). (C) Full-length BicA dimer predicted by AlphaFold2.

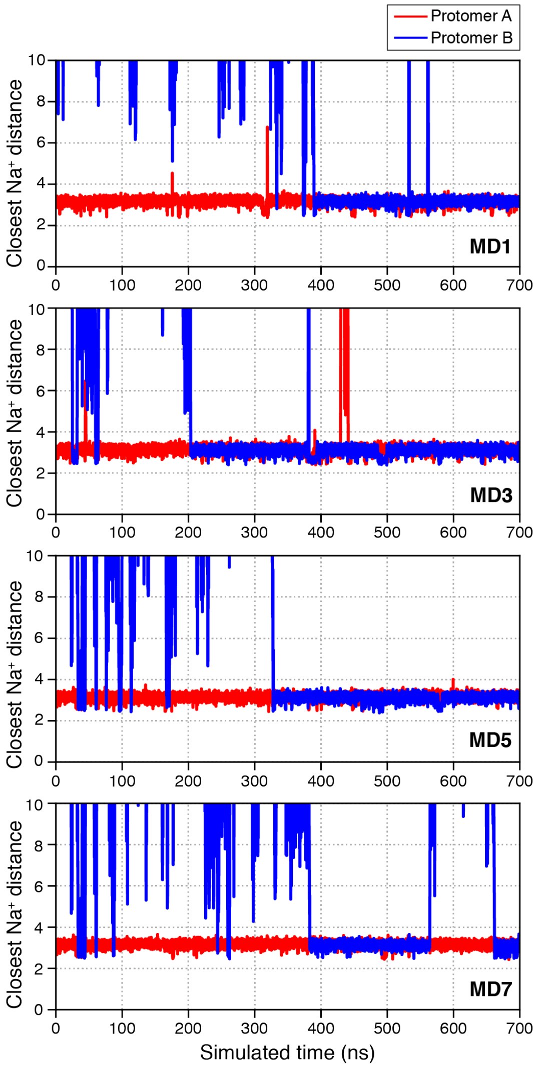


**Figure S15: Binding of the sodium ion to the substrate binding site.** Time-resolved distances measuring the distances of the closest sodium ion to the C$\gamma$ atom of Asp258. A sodium ion bound to BicA protomer A (red line) within the pre-production stages of MD simulation. A total of 7 MD replicates of 700 ns were performed.

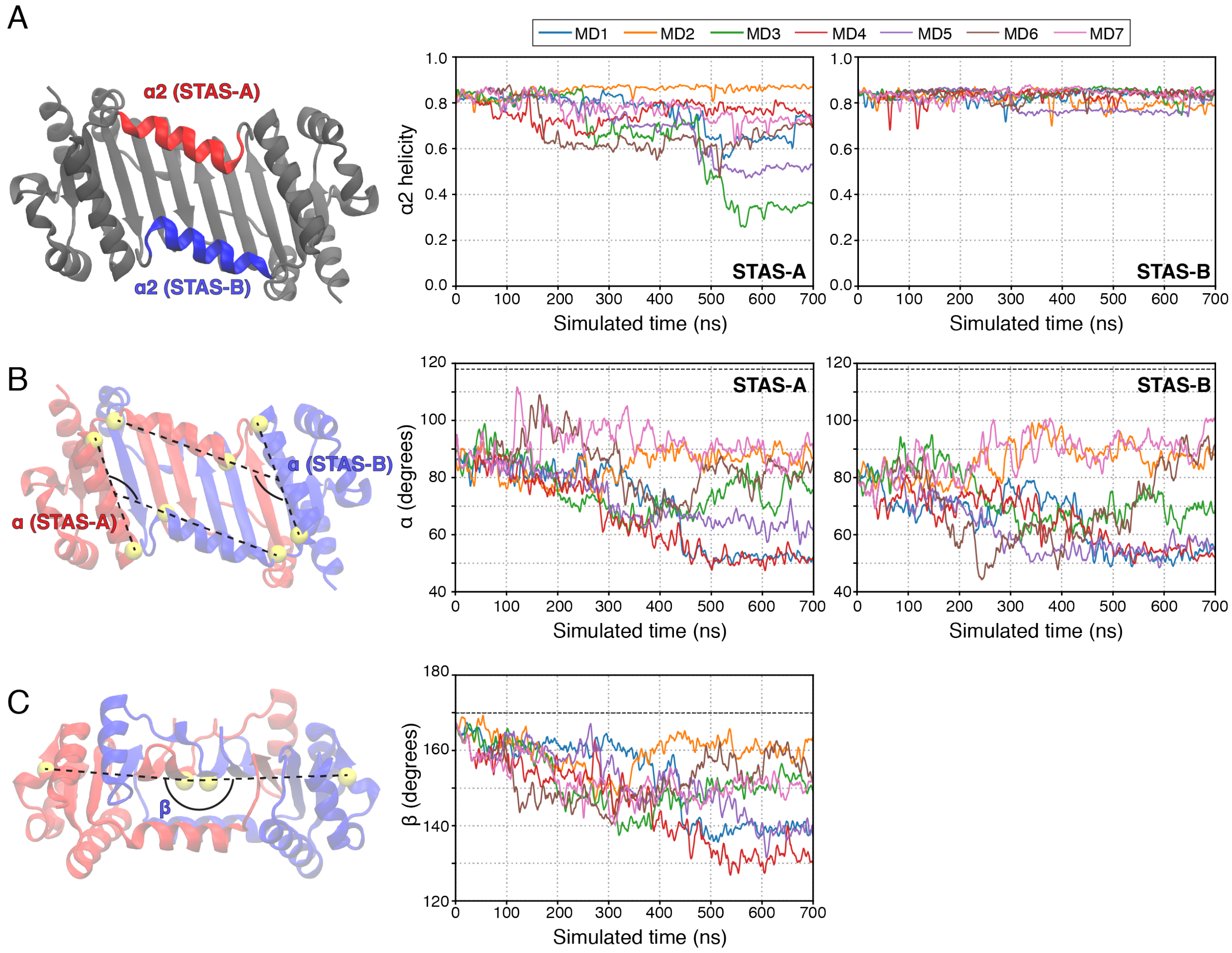


**Figure S16: Structural deformation of the STAS domain.** (A) The helicity of the $\alpha 2$ helix of the STAS domain. Crystal structure of the STAS domain viewed from the intracellular plane. $\alpha 2$ helices are colored in red and blue. Accompanying measurements of helicity over the 7 MD simulations. Individual MD replicates are plotted colored accordingly. (B) Measurement of the angle $\alpha$ formed by $\alpha 2$ and $\alpha 3$ helices, as shown on the STAS structure. Black dashed line indicated the angle observed in the crystal structure. (C) Measurement of the angle $\beta$ formed by the parallel $\beta$ sheets, as shown on the STAS structure. Black dashed line indicated the angle observed in the crystal structure.

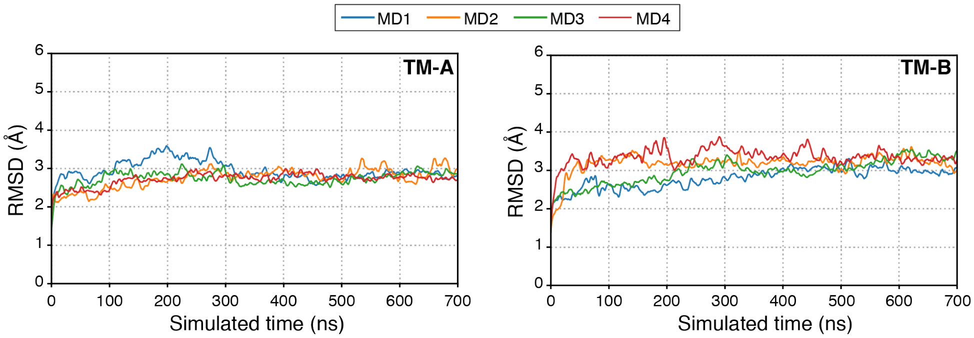


**Figure S17: Stability of the BicA transmembrane domains in simulations in which the STAS domain was not modeled.** Time-resolved root mean squared deviations (RMSD) of individual BicA transmembrane (TM) domains across 4 MD simulations. RMSD was calculated based on the starting structure of the 700 ns simulation. Individual MD replicates are colored accordingly.

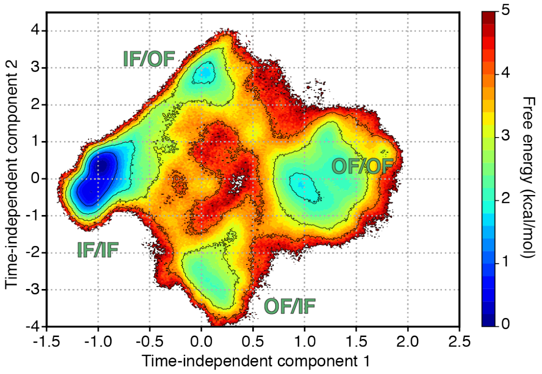


**Figure S18: Projection of the MSM-weighted simulation data on the first two time-independent components calculated by time-independent decomposition.** Time-independent component 1 corresponds to a linear combination of residue-residue distances that contribute to the slowest timescale process. Likewise, time-independent component 2 corresponds to the second slowest timescale process.

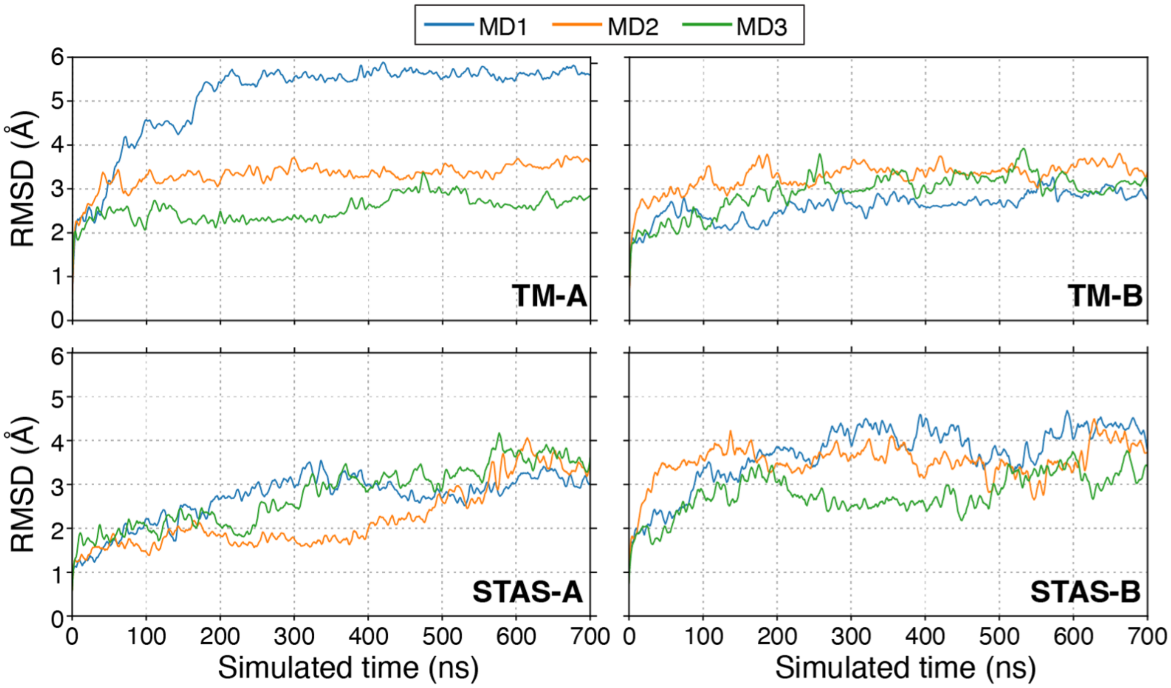


**Figure S19: Stability of the outward facing state seeded from TMD simulations.** Time-resolved root mean squared deviations (RMSD) of individual BicA transmembrane (TM) and STAS domains across 3 MD simulations. RMSD was calculated based on the starting structure of the 700 ns simulation. Individual MD replicates are colored accordingly.

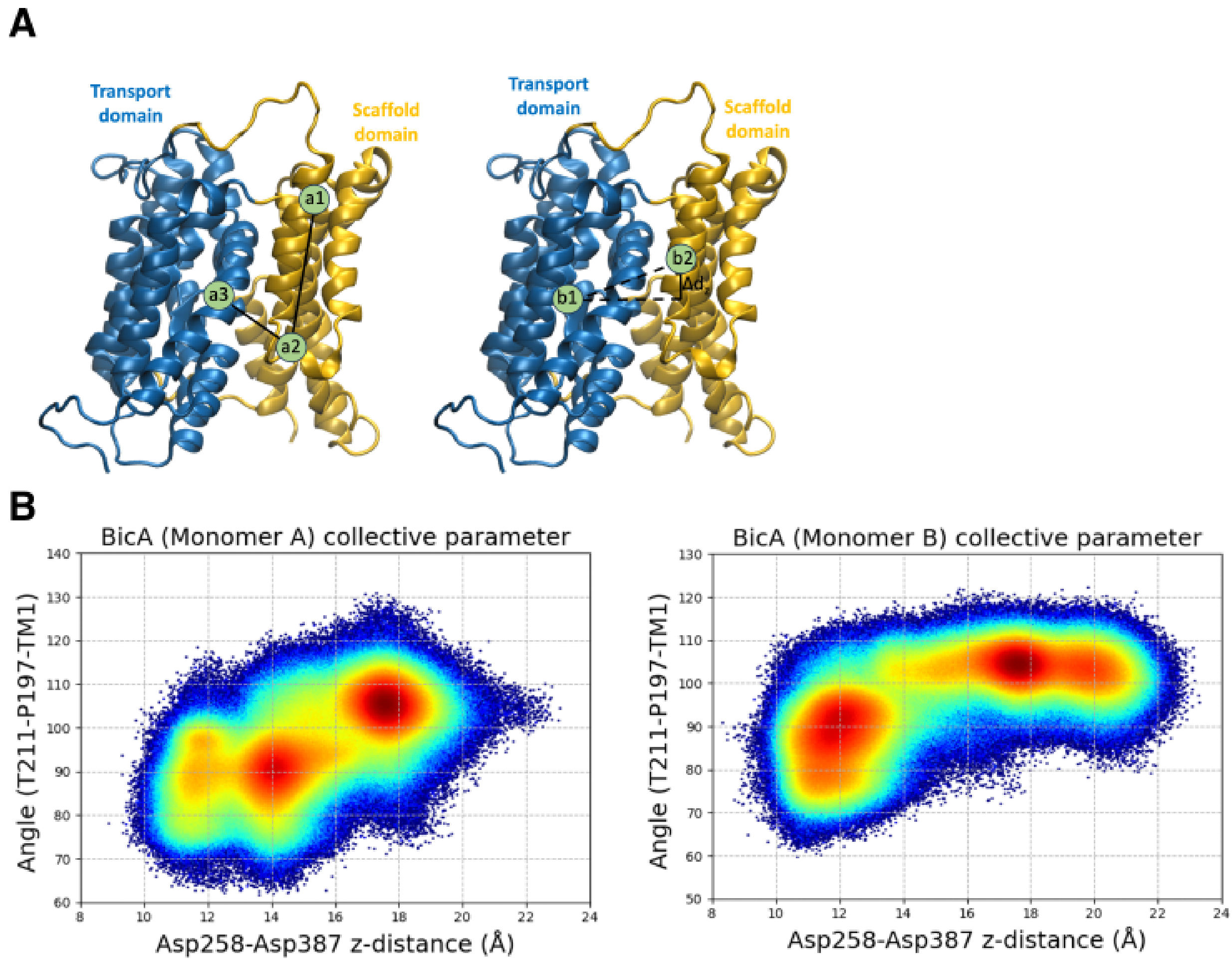


**Figure S20: Projection of the simulation data on angular and translation motion.** (A) Descriptors for angular and translational motion of BicA elevator domain Structure of the transmembrane domain of BicA, colored by transport and scaffold domain in blue and yellow respectively. (Left) Atom a1, a2 and a3 describes the angle for rotational motion of transport domain. Atom a1 (C$\alpha$ atom of T211) and a2 (C$\alpha$ atom of P197) corresponds to beginning and end of TM7 of scaffold domain and atom 3 is the center of mass of TM1 residues (R10-I35), corresponding to the elevator domain. (Right) Representation of the translational motion of elevator domain with respect to scaffold domain is described by the z-axis difference of center of transport domain b1 (C$\alpha$ atom of Asp258) and center of elevator domain of scaffold domain b2 (C$\alpha$ atom of Asp387. (B) Frequency of states define by the rotational and translational motion of the elevator domain. Frequency is colored from low (blue) to high (red).

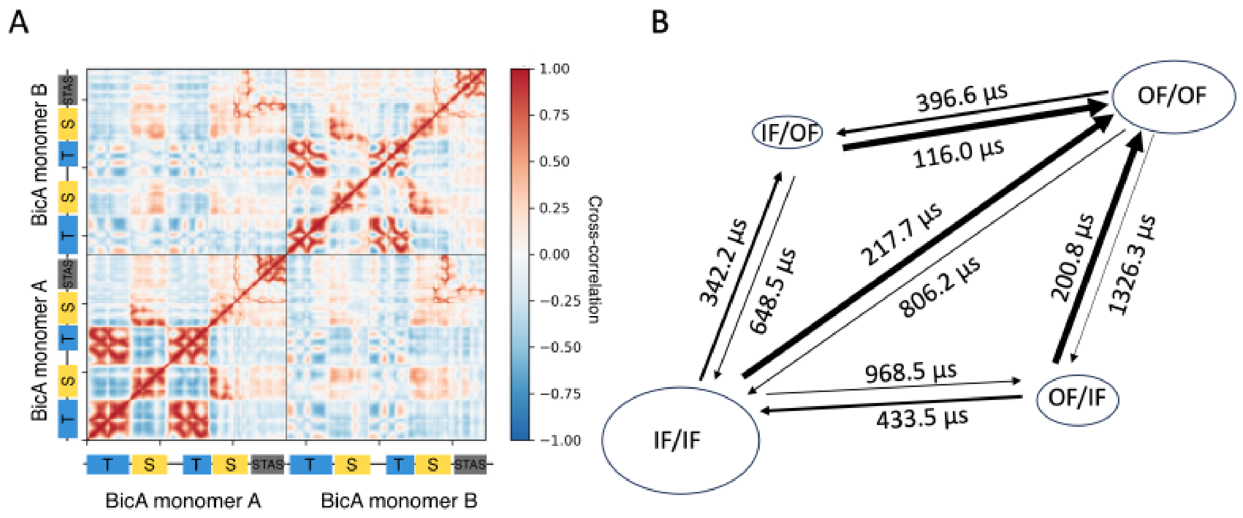


**Figure S21: Independent transitions of BicA protomers** (A) Dynamic cross-correlation motion between the BicA in IF/IF state and BicA in OF/OF state. The two BicA monomers are sequentially ordered on both axes. Transport (T), scaffold (S), and STAS domains are indicated. (B) Mean first passage timescales for conformational transitions between different states of the BicA dimer determined from transition path theory. The arrow thickness represent the relative flux between transitions. The shape sizes surrounding the states correspond to depth of energy minima in Figure4A.

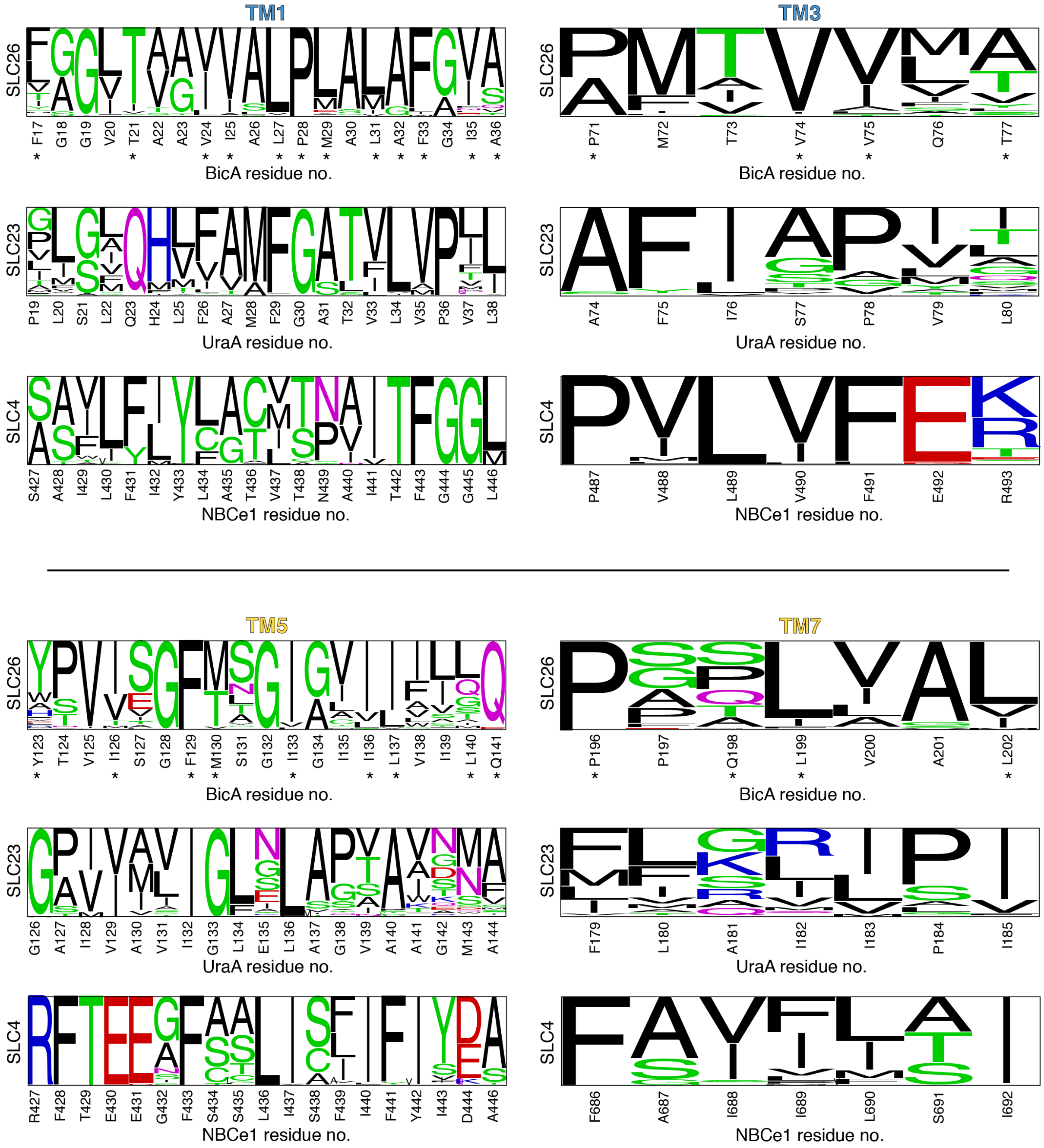


**Figure S22: Amino acid variance among the gating helices of SLC26, SLC23, and SLC4 transporters.** Sequence logo representation of residues on transmembrane helices 1, 3, 5, and 7. Size of the amino acid font represents its respective frequency in the multiple sequence alignment of 300 homologs for the respective transporter family. Residue identified to directly form hydrophobic contacts in BicA MD simulations are marked with (*). The sequence logo of SLC23 and SLC4 homologs are shown for comparison with UraA and NBCe1 sequences used as reference, respectively.

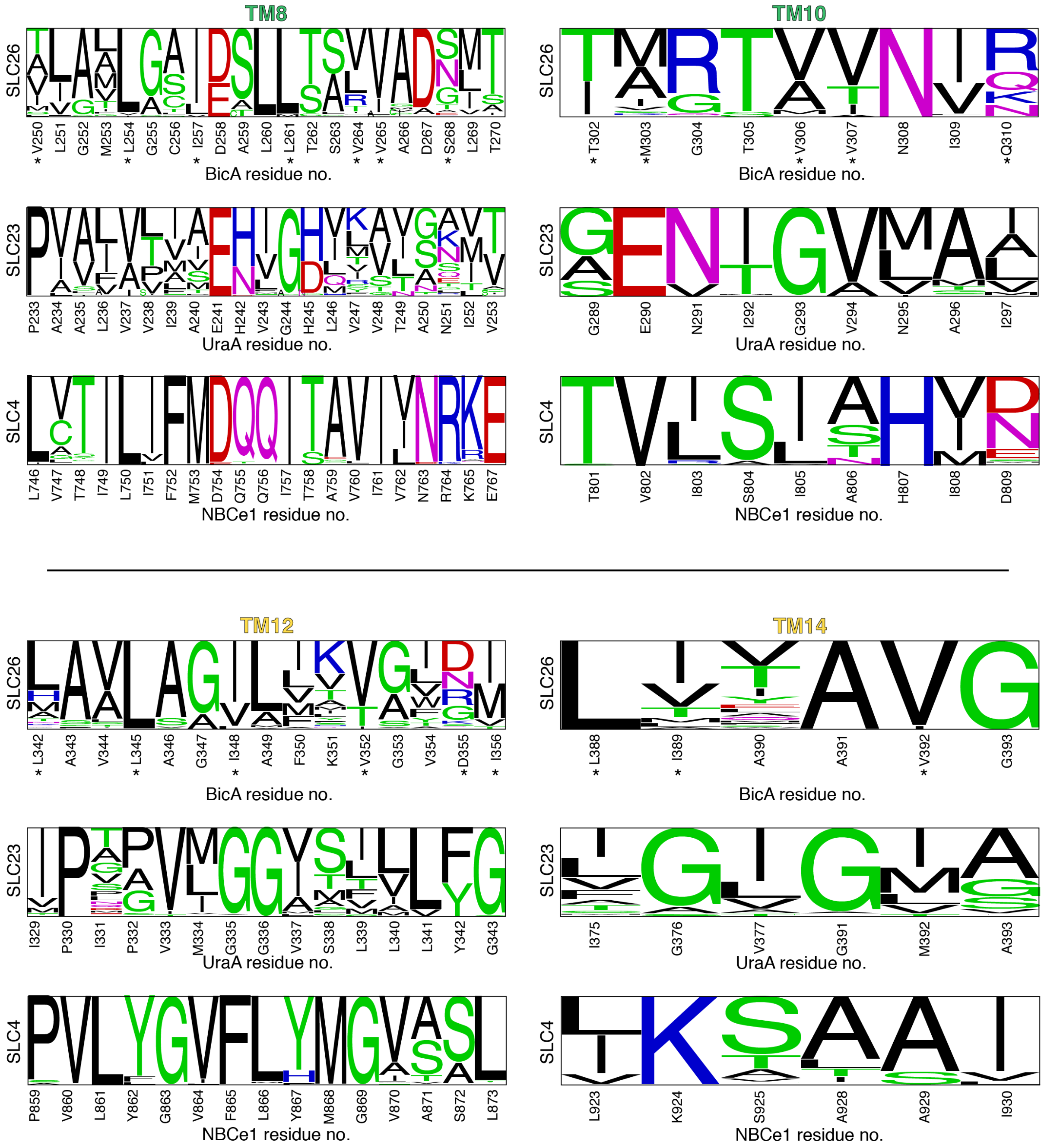


**Figure S23: Amino acid variance among the gating helices of SLC26, SLC23, and SLC4 transporters. Continued from Figure S22.** Sequence logo representation of residues on transmembrane helices 8, 10, 12, and 14. Size of the amino acid font represents its respective frequency in the multiple sequence alignment of 300 homologs for the respective transporter family. Residue identified to directly form hydrophobic contacts in BicA MD simulations are marked with (*). The sequence logo of SLC23 and SLC4 homologs are shown for comparison with UraA and NBCe1 sequences used as reference, respectively.

### Supplemental Tables

**Table S1: Pre-production simulation parameters for the full-length BicA dimer**

| Step | Ensemble | Temperature (K) | Restrained atoms | Force constant (kcal/mol-Å$^2$) | Time (ns) |
|---|---|---|---|---|---|
| 1 | NVT | 10 | All protein atoms | 5 | 2 |
| 2 | NPT | 10 | All protein atoms | 5 | 2 |
| 3 | NPT | 100 | All protein atoms | 5 | 1 |
| 4 | NPT | 200 | All protein atoms | 5 | 1 |
| 5 | NPT | 300 | All protein atoms | 5 | 1 |
| 6 | NPT | 300 | All protein atoms | 5 | 25 |
| 7 | NPT | 300 | Protein backbone | 5 | 25 |
| 8 | NPT | 300 | STAS protein backbone | 5 | 25 |
| 9 | NPT | 300 | STAS C$\alpha$ | 1 | 25 |
| 10 | NPT | 300 | None | N/A | 50 |

**Table S2: Adaptive sampling rounds of full-length BicA dimer simulations**. Individual trajectories were 60 ns in length.

| **Adaptive round** | **Number of trajectories** | **Simulation time ($\mu$s)** |
|---|---|---|
| 1 | 200 | 12.0 |
| 2 | 200 | 12.0 |
| 3 | 200 | 12.0 |
| 4 | 200 | 12.0 |
| 5 | 95 | 5.7 |
| 6 | 95 | 5.7 |
| 7 | 95 | 5.7 |
| 8 | 95 | 5.7 |
| 9 | 100 | 5.8 |
| 10 | 99 | 5.8 |
| 11 | 100 | 6.0 |
| 12 | 100 | 5.9 |
| 13 | 230 | 13.5 |
| 14 | 800 | 47.5 |
| 15 | 625 | 38.4 |
| 16 | 200 | 11.8 |
| 17 | 624 | 39.4 |
| 18 | 600 | 47.9 |
| 19 | 500 | 30.0 |
| 20 | 2997 | 186.7 |
| 21 | 599 | 33.3 |
| 22 | 599 | 37.2 |
| 23 | 599 | 37.3 |
| 24 | 4000 | 266.4 |
| 25 | 400 | 24.0 |
| 26 | 400 | 23.7 |
| 27 | 115 | 6.8 |
| 28 | 441 | 26.0 |
| 29 | 600 | 38.8 |
| Total | | 1002.9 |

**Table S3: Residue distances used for MSM features.** The 96 distances used for one BicA protomer (protomer A) is listed. Residues labeled with "-B" indicated the residue of the other BicA protomer (protomer B). All distances were measure based on C$\alpha$-C$\alpha$ distances.

| Residue 1 | Residue 2 | Residue 1 | Residue 2 | Residue 1 | Residue 2 |
|---|---|---|---|---|---|
| G14 | L317 | G128 | L260 | A470 | R502 |
| L16 | Q310 | G132 | T302 | A470 | D541 |
| V20 | P197 | L140 | L345 | V473 | A489 |
| L27 | P197 | P144 | L176 | F474 | D534 |
| M29 | A43 | P144 | A349 | V504 | S529 |
| A30 | G40 | L146 | I218 | V504 | K540 |
| L31 | V344 | V172 | I218 | T513 | P527 |
| I35 | L140 | M179 | A343 | C531 | S536 |
| A36 | Q76 | I183 | P197 | T119 | N400-B |
| A41 | Q225 | I184 | F350 | D267 | D533-B |
| A41 | A337 | S209 | I218 | N400 | K116-B |
| L45 | G227 | R219 | A334 | L402 | L447-B |
| L57 | G284 | R220 | G332 | R406 | L444-B |
| G59 | H274 | E223 | A334 | S419 | S412-B |
| P62 | L269 | L228 | L292 | S419 | V415-B |
| L64 | M303 | A266 | R271 | N430 | D534-B |
| G70 | M253 | D267 | L427 | R433 | D534-B |
| V75 | G295 | L269 | N308 | G440 | Q467-B |
| S82 | A93 | T270 | R433 | R441 | I472-B |
| A93 | F236 | N308 | G313 | F445 | S477-B |
| A95 | S290 | I309 | V354 | G449 | V484-B |
| F96 | Q232 | L327 | G332 | P450 | L482-B |
| L104 | L251 | K363 | K432 | A460 | L435-B |
| F105 | E279 | A365 | M376 | N464 | G440-B |
| A108 | K113 | H366 | L435 | S477 | F445-B |
| A108 | G255 | I370 | I401 | P480 | G449-B |
| L112 | S276 | I370 | A421 | V484 | R461-B |
| L114 | E273 | T403 | S419 | Y532 | D267-B |
| T119 | F445 | Q411 | R461 | D534 | N430-B |
| P122 | I357 | A421 | E431 | S536 | T270-B |
| P122 | L427 | I452 | A460 | A538 | R433-B |
| G128 | A391 | Q467 | G501 | G547 | T270-B |
